# Internal Dynamics Interact with Proprioceptive Feedback During Movement Execution in an RNN Model of Motor Cortex

**DOI:** 10.1101/2023.12.05.570033

**Authors:** Hongru Jiang, Xiangdong Bu, Zhiyan Zheng, Xiaochuan Pan, Yao Chen

## Abstract

Proprioceptive feedback provides the information about the state of the body, which is critical in motor control. However, the contribution of proprioceptive feedback to motor cortical activity during voluntary movement execution is unclear. Here, we built an recurrent neural network model of motor cortex that receives proprioceptive feedback, and optimized it to control a virtual arm to perform a delayed-reach task. Simulated neural activity is similar to real data, indicating that our model captures the motor cortical dynamics. We further disrupted recurrent connectivity and proprioceptive feedback to dissect their contribution, and found that internal dynamics dominate in neural population activity, while proprioceptive feedback controls movement termination. Moreover, proprioceptive feedback improves the network’s robustness against noisy initial conditions. We further investigated the relative importance of the components in proprioceptive feedback and found that the feedback of hand velocity contributes most to the similarity between simulation and real data. Finally, we show that our motor cortex model can be implemented in the sensorimotor system, demonstrating our model’s biological plausibility. In summary, motor command may arise from the intersection between recurrent dynamics in motor cortex and proprioceptive feedback.

## 1. Introduction

Voluntary movement, such as reaching, grasping, and moving objects, is central to our interaction with the world. The motor cortex (MC), which orchestrates intricate neural pathways to generate motor command, is critical for planning and executing movement. Electrophysiological recordings in non-human primates have revealed that neuronal activity in the MC represents movement kinematics, including direction (Georgopoulos et al., 1982), velocity (Wang et al., 2007) and isometric force (Sergio et al., 2005). However, this perspective only partially explains complex single-neuron multiphasic firing rate (Kalaska, 2019).

With advances in recording technologies and computational methods, the focus on MC has shifted from the representation of output to how motor command is created (Figure 1A). Firing rates of all neurons (neural population responses) in MC coordinate patterns for a particular movement. Recent studies have emphasized that MC is a dynamical system where future state depends on its current state and dynamical rule that dictates evolution (Churchland et al., 2012, 2010; Shenoy et al., 2013; Vyas et al., 2020). The dynamics result from the internal dynamics within MC (neuronal connectivity) and external inputs from other areas. Under the dynamical systems view, Churchland et al. (2012) projected population responses into low-dimensional space by their own dimensionality reduction method named jPCA and found rotational dynamics of the MC when monkeys perform reach task. Neural population responses in the state space from different starting point, follow rotational paths in the same direction for all movements. These rotational patterns revealed the simple nature of complex individual neural responses (Churchland et al., 2012). Furthermore, the dynamical system perspective has paved the way for interpreting motor cortical activity via recurrent neural networks (RNNs) — abstract neural computational of interconnected neurons. In one class of models, the MC is approximated as an autonomous dynamical system during movement execution, with negligible external input (Figure 1B) (Churchland et al., 2012; Hennequin et al., 2014; Sussillo et al., 2015; Zimnik and Churchland, 2021; Michaels et al., 2016). Neural population responses evolve from a specific initial condition, defined as the firing rates at movement onset (Afshar et al., 2011; Churchland et al., 2010), under the control of strong internal dynamics within MC. These models not only generate neural responses similar to real data at singlecell level, but also display rotational dynamics at population level (Churchland et al., 2012). However, the autonomous dynamical system is vulnerable to perturbations in output-relevant dimensions and unable to correct errors from an intended movement. The neural state at movement onset can only vary in a limited subspace, fail tolerate large neural variability.

**Figure 1:**
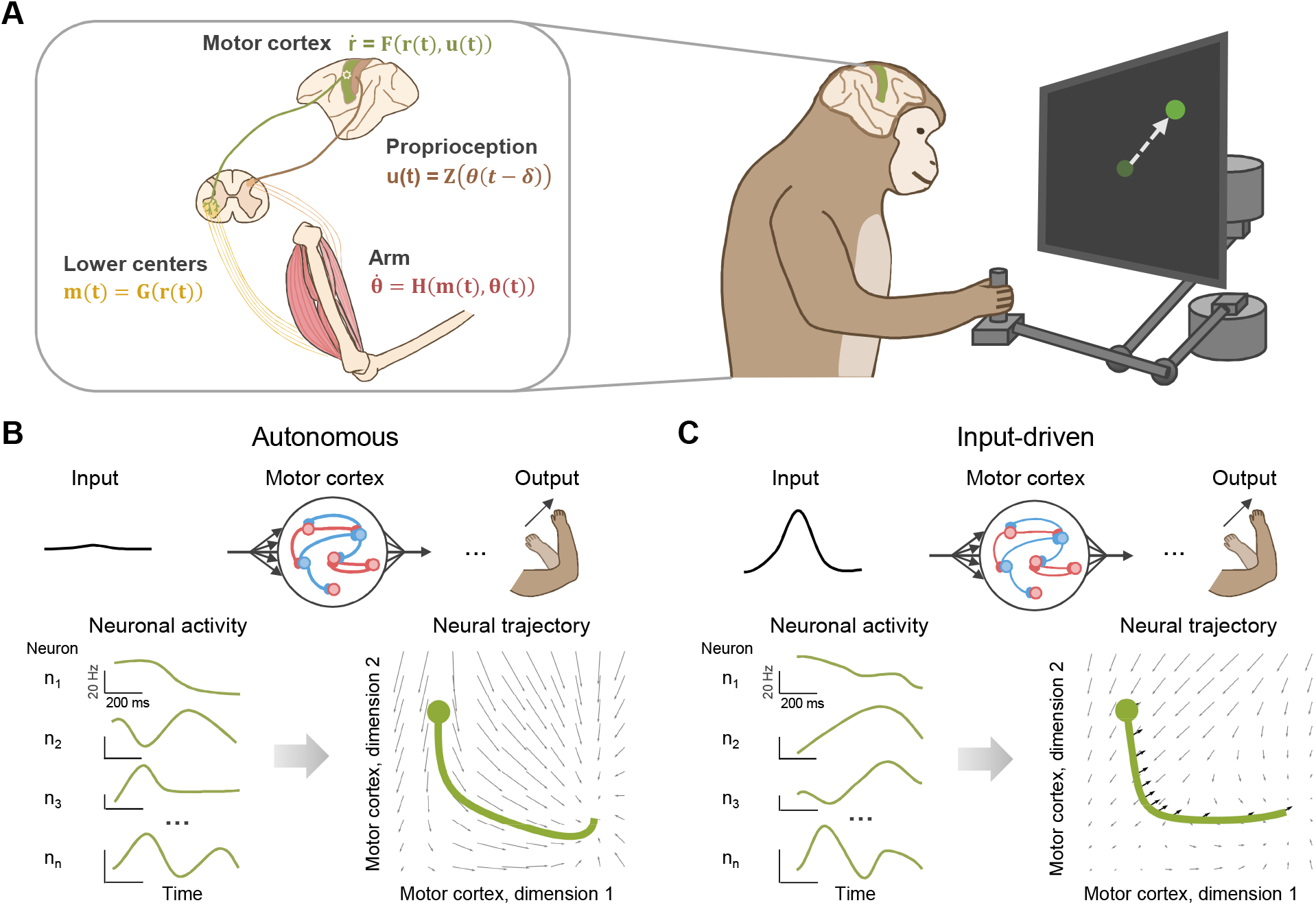
Motor cortex controls reaching under the perspective of dynamical system. (A) Dynamical system perspective on reaching movement. A monkey puts its hand on a planar manipulandum to the center of the workspace to initiate the trial. After a randomized delay, a peripheral target appears, indicating the location the monkey is to reach. Monkey was trained to hold during a variable delay period until a go cue (e.g., auditory signal) prompts the monkey to reach the target. During his reaching, motor cortical firing rates, evolving with a derivative 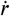, are determined by the internal dynamics in the MC and external input. Firing rates is converted into muscle activity *m*(*t*) via a downstream circuit in lower motor centers, thereby updating the joint angles and velocities θ(*t*), ultimately reflected in hand movement. Delayed sensory feedback from the arm is sent back to the brain regions outside the MC. (B) MC as an autonomous dynamical system during movement execution. Neuronal population responses are visualized in a low-dimensional state-space, with neural states evolving from specific initial conditions (solid circle) under strong internal dynamics within the MC (grey arrows). The length of arrows illustrates the strength of dynamics. (C) MC as an input-driven dynamical system during movement execution. Neural firing rates starting from the initial condition, are driven by external input (black arrows) and internal dynamics (grey arrows). The internal dynamics are weaker than (C).

Alternatively, there is a growing recognition of the role that external inputs play in shaping motor cortical activity (Figure 1C). Situated within a vast and distributed brain network, the MC integrates various external inputs, such as sensory feedback (Sauerbrei et al., 2020; Kalidindi et al., 2021; Gallego et al., 2020), thalamic (Sauerbrei et al., 2020; Logiaco et al., 2021; Yu et al., 2022) and cerebellar signals (Kumar and Ma, 2023). Sensory feedback is critical in finely controlled movements, as the MC has to integrate multiple sensory information including proprioception, vision and tactile sensation, to generate precise commands. The dynamical system perspective is also reflected in the framework of optimal feedback control (OFC), wherein the MC acts as a feedback controller using the best estimate of sensory feedback to produce motor output (Lillicrap and Scott, 2013; Omrani et al., 2017; Todorov and Jordan, 2002). OFC, which has successfully explained experimental behavior (Scott, 2004; Todorov and Jordan, 2002), is also expected to shape motor cortical dynamics during movement execution (Herter et al., 2009; Perich et al., 2020). A recent study found that sensory feedback also exhibits rotational dynamics, suggesting that sensory feedback may be determinant in cortical pattern generation (Kalidindi et al., 2021). Yet for the complete movement process, this model simplifies the external input as merely sensory information during preparatory period, potentially overstating the contribution of sensory feedback during movement execution.

Our study focuses on proprioceptive feedback, which provides the information about the state of the body. The absence of proprioception leads to significant motor deficits, even when visual feedback is available (Sainburg et al., 1995). Healthy individuals estimate hand motion rely almost on proprioceptive feedback compared to visual feedback (Crevecoeur et al., 2016). Utilizing the inhibitory stabilized network (ISN) (Hennequin et al., 2014; Kao et al., 2021), we built a model of MC that receives the feedback of hand position and velocity, and muscle force. Our model generated firing rates that resembled real data at both single-cell and population levels, both qualitatively and quantitatively. We found that both recurrent structure and proprioceptive feedback are important in motor cortical dynamics. To dissect their contribution, we disrupted them at graded percentage of timecourse. The absence of internal dynamics causes neural activity out of control and significant deviation of the target, whereas the absence of proprioceptive feedback led to ceaseless movement. These results suggest that internal dynamics are dominant in cortical pattern generation, while proprioceptive feedback is essential for velocity control. In an initial condition perturbation experiment, proprioceptive feedback improved the network robustness. We further investigate which component in proprioceptive feedback is important, and found that hand position and velocity is important in model’s robustness, whereas hand velocity is critical in motor cortical dynamics. Finally, we implemented our MC model can be in the sensorimotor system, demonstrating our model’s biological plausibility.

In summary, our research offers new insights into the intricate interaction between the internal dynamics within MC and proprioceptive feedback during movement execution, potentially reshaping our understanding of motor control.

## 2. Method

In this section, we describe the model design for reach generation and methods to evaluate and analyze model. We first describe the building blocks of our model, and explains the loss function calculation and training methods. Finally, we introduce comparison of the model to real data and analyzing its population dynamics.

### 2.1. Model description

We constructed an RNN of motor cortex to control the arm model (Figure 2). The simulation was divided into two epochs by the gated action of basal ganglia: the preparation-epoch (before go cue) where basal ganglia disinhibited thalamic neurons, leaving them to interact with cortex. The execution-epoch (after go cue) where basal ganglia inhibited thalamus and opened the loop, and movement-specific input is withdraw. The dynamics of MC model evolve from movement-specific initial condition, under the control of internal dynamics and proprioceptive input (composed of task input and proprioceptive feedback), generate firing rate and further muscle activity to actuate arm model. proprioceptive feedback from the arm model is sent back to the motor cortex, closing the loop. Here, we first describe the motor cortex model, inputs and initial condition setup. Then we introduce the arm model and proprioceptive system.

**Figure 2:**
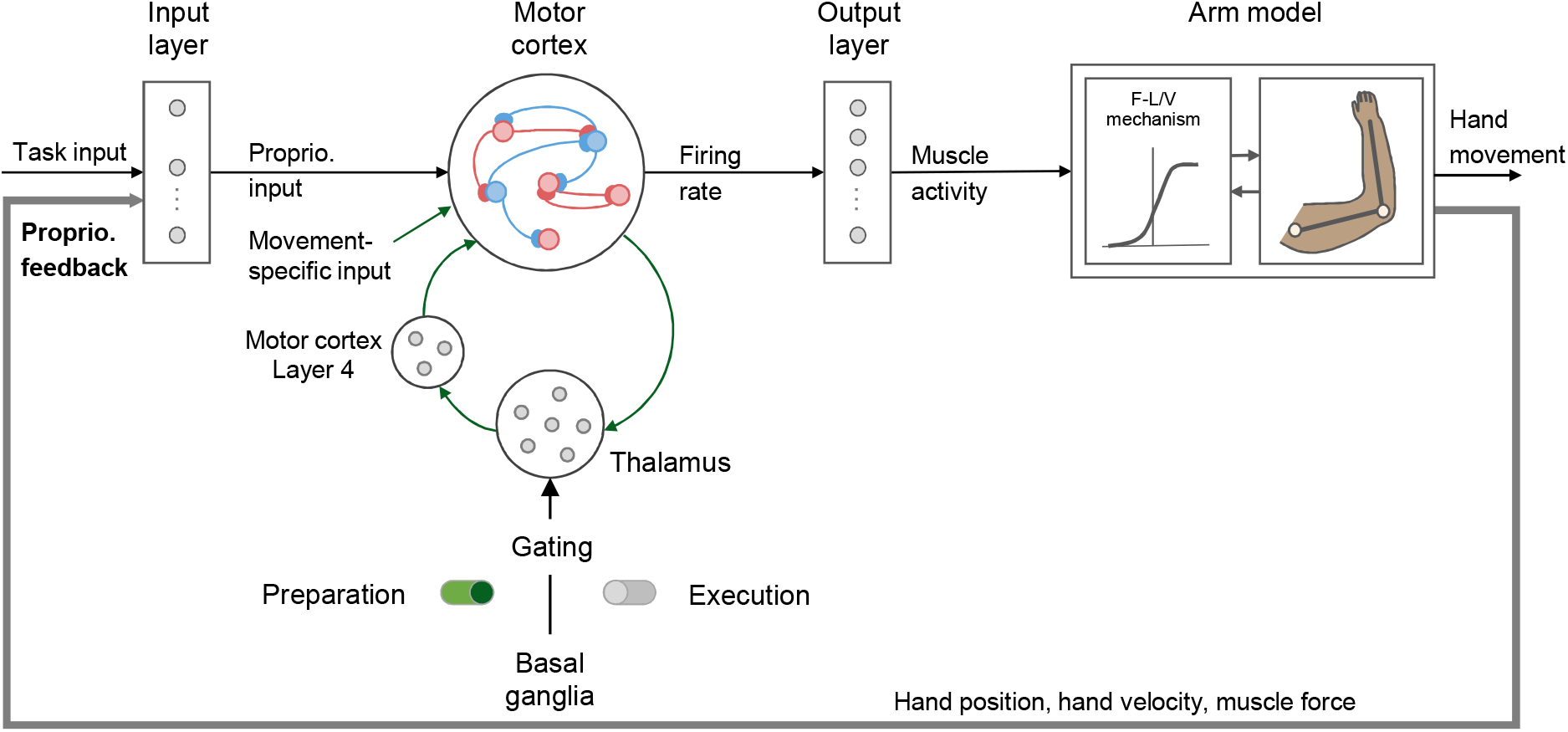
Schematics of modeling framework. The firing rates of MC network are translated into muscle activity via the output layer. Then, the noisy muscle activity is converted into muscle force through F-L/V mechanism, actuating the motion of the two-link arm model. MC receives task and proprioceptive inputs during movement execution. Proprioceptive feedback (hand position and velocity, and muscle force) from the arm ascends into MC. The thalamocortical loop only works during preparation epoch and re-opens after go cue.

#### 2.1.1. Motor cortex model

We modeled the MC (Figure 2) as an biologically plausible network, inhibition-stabilized network (ISN), satisfying Dale’s law (neurons are either excitatory or inhibitory) (Hennequin et al., 2014; Eskikand et al., 2023). Our ISN comprised 300 excitatory and 100 inhibitory neurons (*N* = 400), with a similar neuron proportion to that in the MC of non-human primates (Bakken et al., 2021; Chen et al., 2023). We constructed a random connectivity matrix of size *N* × *N* with a connectivity density *p* = 0.1. The non-zero excitatory weights were drawn from a lognormal distribution with mean *log*(*w*_0_) and variance 1, where 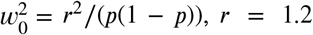. For inhibitory weights, we employed a uniform distribution between −1 and 0. The random excitatory weights are stabilized by fine-tuned inhibitory synapses (Hennequin et al., 2014), which were optimized by minimizing the “smoothed spectral abscissa”, the greatest real part of ***W***’s eigenspectrum, to below 0.9, following the method in Hennequin et al. (2014).

The neuronal activity ***x***(*t*) = *x*_1_(*t*), …, *x*_*N*_ (*t*) ^T^ evolves according to:

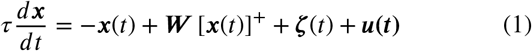

where τ = 200 is the time constant of MC dynamics. []^+^ is the ReLU Activation Function. **ζ** (*t*) = ***z***_*sp*_ + ***z***_*ci*_(*t*) includes two terms: an input ***z***_*sp*_ = ***x***_*sp*_ − ***W*** [***x***_*sp*_]^+^ maintains the spontaneous activity of the network, ***x***_***sp***_, which is sampled from Gaussian distribution 𝒩 (20, 9); a condition-independent signal ***z***_*ci*_, which appears after the go cue and induces a substantial change in dynamics (Kaufman et al., 2016; Kao et al., 2021):

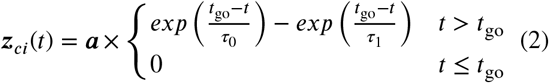

where ***a*** = 7.28 + **ζ**. **ζ** is a N dimensional vector sampled from Gaussian distribution 𝒩 (0, 0.2). τ_0_ and τ_1_ are in table 1. ***u***(*t*) is the external input specified below.

**Table 1.** Parameters of the musculoskeletal arm model.

| Symbol | Description | Value | Unit |
| --- | --- | --- | --- |
| $p_1^{init}$ | initial hand position (x-axixs) | 0 | m |
| $p_2^{init}$ | initial hand position (y-axixs) | 0.4 | m |
| $\theta_1^{init}$ | initial elbow angle | 0.63 | rads |
| $\theta_2^{init}$ | initial shoulder angle | 1.77 | rads |
| $L_1$ | length of upper arm | 0.30 | m |
| $L_2$ | length of lower arm | 0.33 | m |
| $M_2$ | mass of lower arm | 1 | kg |
| $D_2$ | center of mass of lower arm | 16 | cm |
| $I_1$ | moment of inertia of upper arm | 0.025 | kg $m^2$ |
| $I_2$ | moment of inertia of lower arm | 0.045 | kg $m^2$ |
| $\theta_k^{targ}$ | target angle ($k = 1, \dots, 8$) | $45 \times (3 + k)$ | degree |
| $d$ | target distance | 15 | cm |
| $p_1^{targ}$ | relative target position (x-axixs) | $d \times \cos(\theta^{targ})$ | cm |
| $p_2^{targ}$ | relative target position (y-axixs) | $d \times \sin(\theta^{targ})$ | cm |

The readout of MC is muscle activity that actuates musculoskeletal arm model. The muscle activity is defined as:

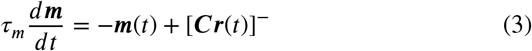

where the readout weight matrix ***C*** is sampled from Gaussian distribution 𝒩 (0, 0.1^2^/*N*), and τ_m_ = 50 ms is the time constant. []^−^ is the LeakyReLU Activation Function with negative slope equal to 0.4.

#### 2.1.2. Inputs of proprioception and preparatory control to the motor cortex

The inputs to MC model, 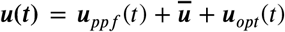, include three terms: a proprioceptive feedback input ***u***_*ppf*_ (*t*), a task input 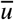 denoting target position, and a preparatory control input ***u***_*opt*_(*t*) (Figure 2). First, proprioceptive feedback is a ten-dimensional vector ***I***_*p*_(*t*) consists of hand position ***P***, hand velocity 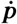, and muscle force ***f***:

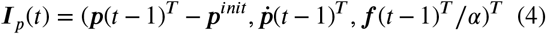

where 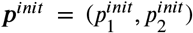 is the initial hand position (see Table 1). α = 2.7 is hyper-parameters. The proprioceptive feedback input to the MC model is:

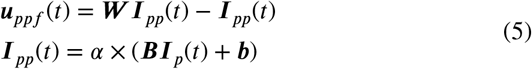

where ***B*** is an *N* ×10 weight matrix sampled from Gaussian distribution 𝒩 (0, 1/10). ***b*** is the bias, initialized to 0.

Second, the task input informs the MC where the target is, defined as:

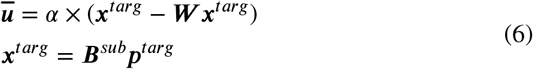

where ***B***^*sub*^ is the first two columns of 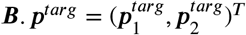 is the target position relative to the the origin (see Table 1). The above definitions make MC network compared the current hand position to the target position (explained in the appendix).

Finally, the preparatory control input is the combination of thalamic inputs and a movement-specific input, and switched off at go cue onset:

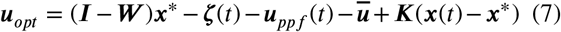

where ***x***^∗^ is the initial condition specified in next section. ***K*** is the optimal feedback matrix and calculated from the method described in (Kao et al., 2021). The dynamics of gated thalamocortical circuit model are in appendix.

#### 2.1.3. Initial condition of motor cortex based on proprioceptive input

Neural responses in MC shows an rich transients after go cue onset (Churchland et al., 2012; Shenoy et al., 2013). As the target has been present until the end of the movement, the task input should be consisten after target onset. To detach the contribution of task input in setting initial condition, we defined the movement-specific motor cortical state at go cue onset as the initial condition of MC model, 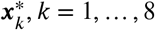,as:

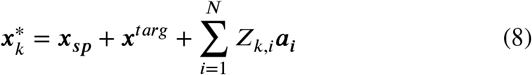

where ***a*** = (***a***_**1**_, ***a***_**2**_, …, ***a***_***N***_)^*T*^ is the full eigenvectors of ***O***, ordered by decreasing eigenvalues, and ***Z*** is a weight matrix sampled from Gaussian distribution 𝒩 (0, 1.5/*N*). The initial conditions contains the information about spontaneous activity, target position and various transient behaviors. We also scaled the variance of firing rates after go cue to 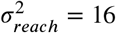 Hz (Kao et al., 2021).

#### 2.4.1. Arm model

The two-link musculoskeletal arm model (Figure 2) is in the horizontal two-dimensional plane, driven by six lumped muscles (mono-articular shoulder flexors, mono-articular shoulder extensors, mono articular elbow flexors, monoarticular elbow extensors, biarticular flexors and biarticular extensors) (Lillicrap and Scott, 2013). First, muscle activity ***m***(*t*) was transformed into muscle force ***f*** (*t*) via the forcelength-velocity function:

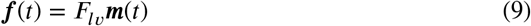

where 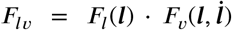 is the force-length-velocity function (the parameters are in appendix):

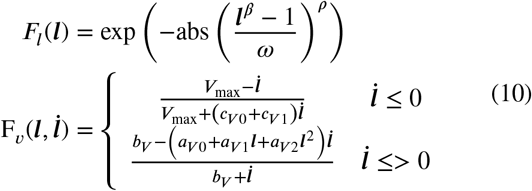

where ***l*** = (*l*_1_, *l*_2_, …, *l*_6_) and 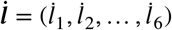 are the length and velocity of each muscle, defined as:

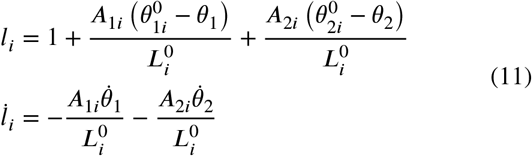

where 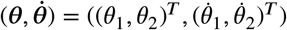 includes the angles and velocities of the elbow and shoulder joints, which are initialized with the parameters in Table 1 at the trial beginning. *A* is the moment arm matrix, ***L***^**0**^ is the optimal length matrix and **θ**^**0**^ are the and joint angle. These parameters are defined in the appendix.

The muscle force was mapped into joint torques with a delay of δ_*f*_ = 100 ms, as the time when muscle activity begin to change lead movement onset in the experimental observations (Sussillo et al., 2015):

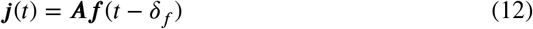

The joint kinematics evolved dynamically according to:

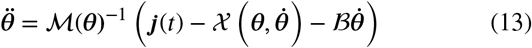

where ℳ is the inertia matrix, 𝒳 is the centripetal and Coriolis forces, and ℬ is the damping matrix representing joint friction. The above parameters are given by

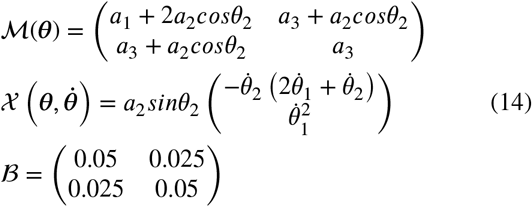

where 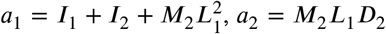 and *a*_3_ = *I*_2_. These parameters are given in the Table 1. The hand position, ***p*** and velocity, 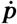 are given by:

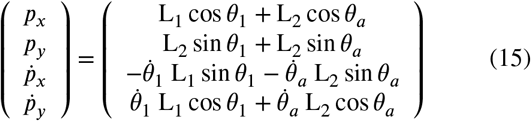

where 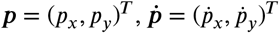, and θ_*a*_ = θ_1_ + θ_2_.

#### 2.1.5. Proprioceptive system model

We specified the proprioception pathway into a proprioceptive system model, and then incorporated with our motor cortex model for more complete simulation of sensorimotor system.

The representation of proprioception has been widelt modeled in the long short-term memory (LSTM) network (Vargas et al., 2024). Here, we utilized recurrent unit (GRU), a streamlined version of the LSTM that often achieves comparable performance but faster computation (Chung et al., 2014). In order to estimate the proprioception from the noisy and delayed proprioceptive feedback and the efference copy of the motor command (muscle activity). The input is defined as:

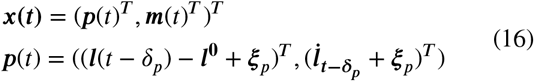

where *l* is the muscle length 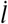 is muscle velocity, and *l*^0^ is the muscle length at the trial beginning. δ_*p*_ = 35 ms, as the proprioceptive feedback takes at least 35 ms to reflect on the motor cortical population activity (Pruszynski et al., 2014). **ξ**_*p*_ is the sensory noise (specified in 3.5). The details of computation are in appendix.

### 2.2. Model optimization design

#### 2.2.1. Design of loss function for motor cortex model

In delayed-reach task, the network was required to control the arm model to reach the target within 1000 ms after go cue. For fast computation speed, We first built a baseline model by Euler integration with time steps of *dt* = 10 ms. The model was trained for 1000 iterations by Adam optimizer with a learning rate of 2 × 10^−4^. Before optimization, the initial conditions x^∗^ and spontaneous activity *x*_*sp*_ were restricted in the nullspace of ***C*** to prevent movement from occurring at the end of preparation and spontaneously. The loss function are composed of the MSE between the network output and the target at the end of the movement, and penalization terms:

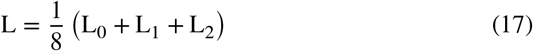

First, the network should hold the arm model for a T_0_ = 300 ms after reaching the target, where hand velocity is nearly zero.

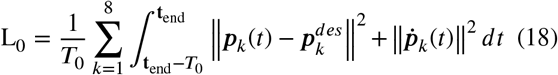

where *k* is the number of movement condition, ***p***^*des*^ = ***p***^*init*^ + ***p***^*targ*^ is the target position, and t_end_ = 1200 ms. We also penalized muscle activity to prevent undue muscle output at the end of the movement:

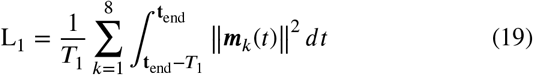

where T_1_ = 200 ms.

Second, we regularized the neural dynamics in a lowdimensional space. We reduce the model and monkey data to ten dimensions by PCA and sample model data every three time points to have the same time length with monkey data. The penalization term is:

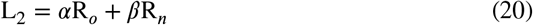

where α = 9 × 10^™3^, β = 10^™4^.

The first term penalizes the shapes of the first two principle components (PCs), as we observed that the first two PCs of real data are generally α-shaped:

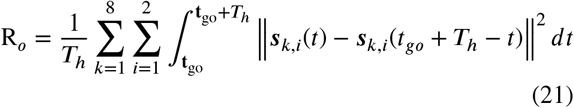

where ***s*** ∈ ***R***^8×10×*t*^ is the PCs, where *t* = 840 is the number of time points, *c* = 8 *T*_*h*_ = 420 ms.

The second term penalizes the temporal distribution of the sixth to the tenth principle components (PCs). If there was no penalization, these PCs would quickly decay to zero after movement onset. We required that the value of these PCs at the first half of movement was no stronger than that of the latter half:

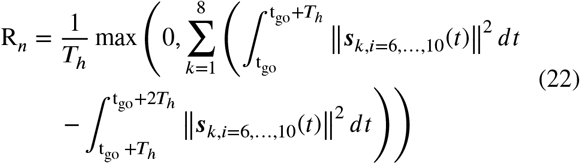

We optimized movement-specific initial condition ***x***^∗^, input weight matrix ***B*** and readout matrix ***C***.

After training, we set *dt* = 1 ms and optimized ***x***^∗^, using the Equation 2.2.1 as loss function.

#### 2.2.2. Design of loss function for proprioceptive system model

We replaced the direct feedback by the proprioceptive system model. First, the proprioceptive system model was trained for 1000 iterations with the Adam optimizer, with a learning rate of 5 × 10^−^4. The loss function was the vector norm between the estimated and the actual proprioception:

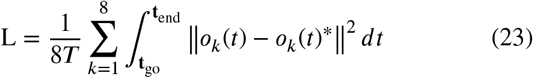

where ***o***_*k*_(*t*)^∗^ is composed of the actual hand velocities and muscle force. The set of parameters to be optimized is (***W***_***rx***_, ***W***_***ux***_, ***W***_***rh***_, ***W***_***uh***_, ***W***_***hx***_, ***W***_***hh***_, ***W***_***oh***_, ***b***_***r***_, ***b***_***u***_, ***b***_***h***_, ***b***_***o***_). Sensory noise was not present during training.

Subsequently, motor noise was added to muscle activity, while sensory noise was added to the proprioceptive feedback:

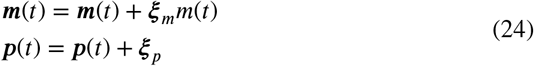

where **ξ**_*m*_ and **ξ**_*p*_ were randomly sampled from Gaussian distribution 𝒩(0, 0.05). The loss function is defined as:

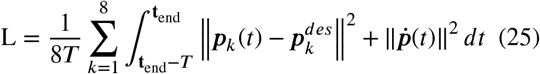

where *T* = 1200 ms and *t*_*end*_ = 300 ms.

Only the initial conditions was optimized and restricted in the nullspace of the readout matrix ***C*** and scaled the variance of firing rates during training. We used the Adam optimizer with a learning rate of 3 × 10^−^3 and 40 iterations.

### 2.3. Quantification and statistical analysis

#### 2.3.1. Preprocessing and Principal component analysis

Neural recording were made from the primary motor cortex (M1) and dorsal premotor cortex (PMd) of Macaque monkey C and M using Utah electrode arrays (Gallego et al., 2020; Gallego-Carracedo et al., 2022; Safaie et al., 2023). Here, We analysed one session of the recordings (268 neurons for monkey C, and 98 neurons for monkey M) and obtained the firing rates following the same procedure described in Safaie et al. (2023). We applied a Gaussian kernel (σ = 50 ms) to the binned square-root transformed spike counts (bin size 30 ms) of each neuron, and excluded neurons whose mean firing rate less than 1 Hz. Firing rates were computed separately for the movement preparation and execution periods: a 1000 ms window starting 200 ms before target onset, and a 1000 ms window starting 300 ms before movement onset.

The movement onset in our model is defined as time when hand velocity begins to change, however, this subtle change is difficult to detect in the experiment. In the analysis of execution period, We set a 1000 ms window starting 250 (150) ms before movement onset (go cue), leaving 50 ms for the detectable changes of hand velocity. Considering the external stimuli takes at time to influence cortical responses and trial variability, we select a period starting *t*_*r*_ (drawn i.i.d. from 𝒩 (150, 20)) before go cue. We perform such temporal alignment when comparing model and real data.

We produced a data matrix ***X*** ∈ ***R***^n×*ct*^, where n is the number of neurons, c is the number of reach conditions, and t is the number of time points. Then we ‘soft’ normalized each neuron’s firing rate (fr): *f r* = *f r*/(*range*(*f r*)+5), and mean-centered fr so that each neuron has a zero mean (Churchland et al., 2012). We used Principal component analysis (PCA) to reduce the dimensionality of data ***X***_***red***_ ∈ ***R***^n×*ct*^, where k is the number of principal components (PCs).

#### 2.3.2. Similarity between model and real data

We used demixed principal component analysis (dPCA) (Brendel et al., 2011; Kobak et al., 2016) to compare key features of the simulated and real population responses, and quantify their similarity by orthogonal canonical-correlation analysis (orthogonal CCA) and Procrustes analysis.

dPCA seeks to find a transformation matrix ***W***, which projects the data matrix ***R***, onto a lower-dimensional space ***X*** through the equation that ***X*** = ***RW***, where each row of ***R*** indicates the condition and time points of firing rates, each column of ***W*** represents a new dimension, and each column of ***X*** represents a component of the population response. The key difference between dPCA and PCA is that dPCA seeks a transformation that not only captures variance but also aligns the principal components with the conditions or time indicated by the labels. Therefore, dPCA found some components (columns of ***X***) varying over time while being constant across conditions, and others varying across conditions while staying constant over time. We divided these two components into: condition-independent and condition-dependent. We also ignored noise covariance within trials.

Orthogonal CCA (Cunningham and Ghahramani, 2015; Russo et al., 2018) is an extension of the traditional CCA that enforces an orthogonality constraint on the transformations, leading to uncorrelated components within each dataset. The similarity between mean-centered datasets ***X***_***a***_ ∈ ***R***^*k*×*ct*^ and ***X***_***b***_ ∈ ***R***^*k*×*ct*^ is:

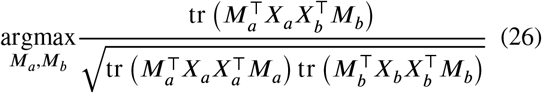

where *M*_*a*_ and *M*_*b*_ are orthonormal matrices. Similarity will be unity if two datasets are the same. Here, we reduce the dimensions by PCA (k=10), which ensures that correlations were not driven by low-variance data (Sussillo et al., 2015). Finally, we compute the similarity.

Procrustes analysis compares two sets of shape data by minimizing the sum of squared differences between corresponding points in the two sets. The method involves scaling, rotating, and translating one set of data to optimally superimpose it onto the other set. We used Procrustes analysis to compare the PCs of model and real data. To assess the goodness of fit, we calculated the *M*^2^, which is the sum of the squares of the point-wise differences between the two datasets. *M*^2^ is 0 indicates a perfect fit, while 1 indicates a poorer fit.

Here, we focused on the same execution period described in 2.3.1. As the time points of monkeys’ firing rates are too few to be applied dPCA, we interpolated the monkey data every three time points by cubic interpolation.

#### 2.3.3. Population ratio

As illustrated in Figure 4B, C, the population ratio quantifies the changes in population activity after different disruptions, defined as:

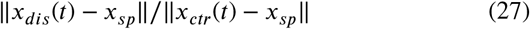

where *x*_*dis*_(*t*) is the neuronal activity in disruption trial, *x*_*ctr*_(*t*) is the neuronal activity in control trial. We first removed the spontaneous activity and computed the magnitude of the population activity, which is the vector norm across all neurons. We then compared this magnitude to that in control trial, and averaged the ratio across all conditions.

#### 2.3.4. Specific direction of initial condition perturbation

In 3.3, the ‘dynamic-direction’ that evokes the most energy variation (Hennequin et al., 2012) of the network dynamics is the first eigenvector of the observability Gramians:

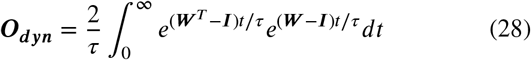

where ***I*** is the unit matrix. ***O***_***dyn***_ can be solved from Lyapunov equation, ***A***^*T*^ ***Q*** + ***QA*** + ***I***^*T*^ ***I*** = 0.

The ‘output-direction’ that evokes the most energy variation of the readout (Kao et al., 2021) is the first eigenvector of the modified observability Gramians:

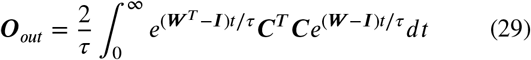

where ***C*** is the readout weight matrix. ***O***_***out***_ can also be solved from Lyapunov equation, ***A***^*T*^ ***Q*** + ***QA*** + ***C***^*T*^ ***C*** = 0.

## 3. Results

### 3.1. Comparison of the model to real data

The delay-reach task is divided into movement preparation and execution by go cue. Our optimized MC network model was able to withhold the arm’s movement until the go cue onset and move the hand to the target position (Figure 3A). The firing rates of individual neurons separated across movement conditions were similar to those observed in monkey data: some neurons display homogeneous, while others display multiphase and heterogeneous firing pattern after movement onset (Figure 3B).

**Figure 3:**
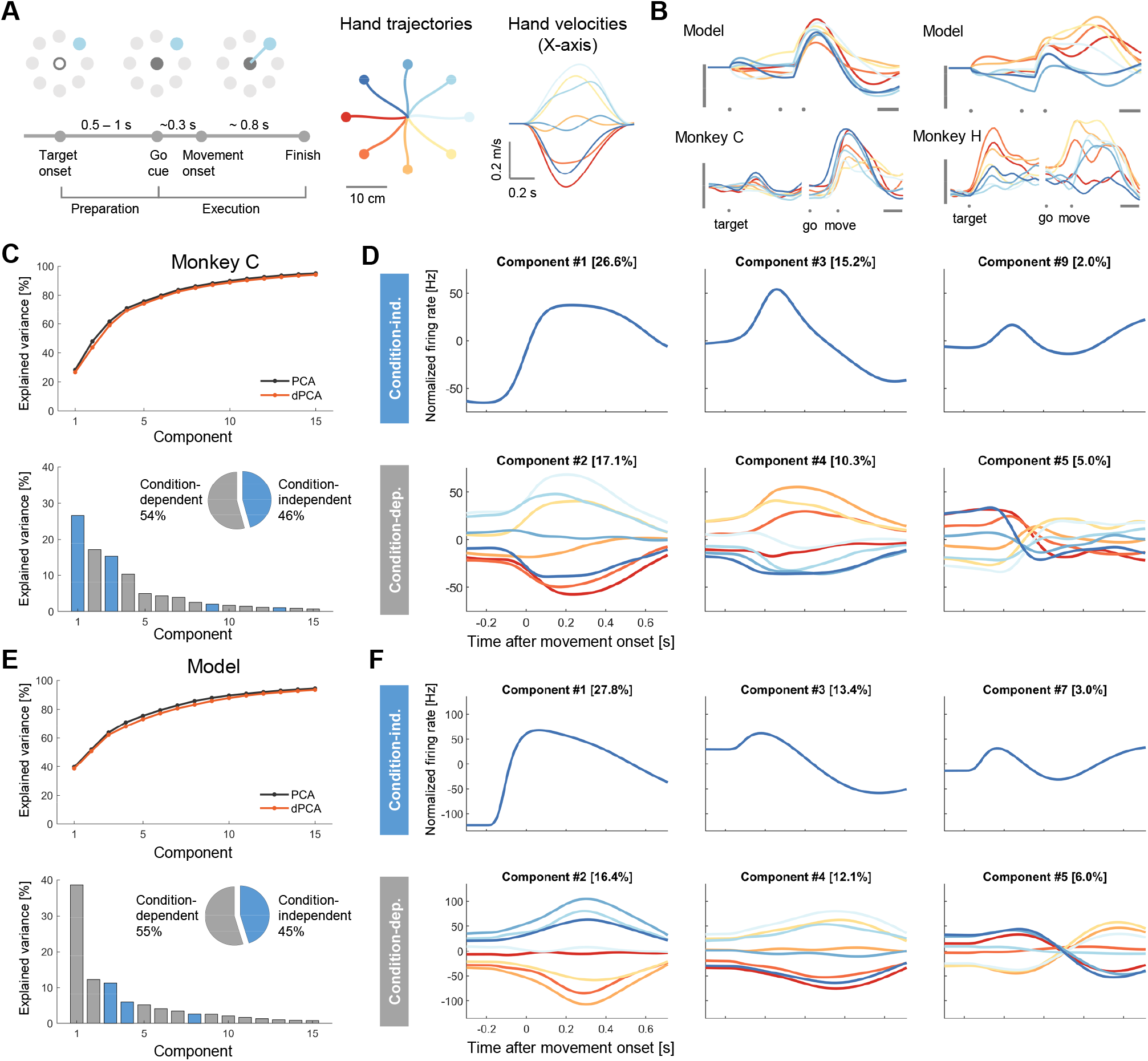
Neural population dynamics of monkey and model during reaching. (A) The simulation of the delayed reach task. Left: the schematic of the task indicating important time point and period for analysis. Middle and right: hand trajectories and velocities generated by the model’s arm. (B) Firing rates of representative neurons in the model and monkey. Traces are colored according to the hand trajectories. Vertical bar: 20Hz, and Horizontal bar: 200 ms. (C) Variance captured by dPCA components for monkey C data. Top: cumulative variance explained by dPCA (black) and PCA (orange). Bottom: bars show the variance of the individual demixed principal components. Gray for condition-independent signal, and blue for condition-dependent signal. Pie chart shows the proportions of each signal. (D) Demixed principal components for monkey C data. Top: first three components for condition-independent signal. Bottom: first three components for condition-dependent signal. (E, F) Same as (C, D) except for model data.

**Figure 4:**
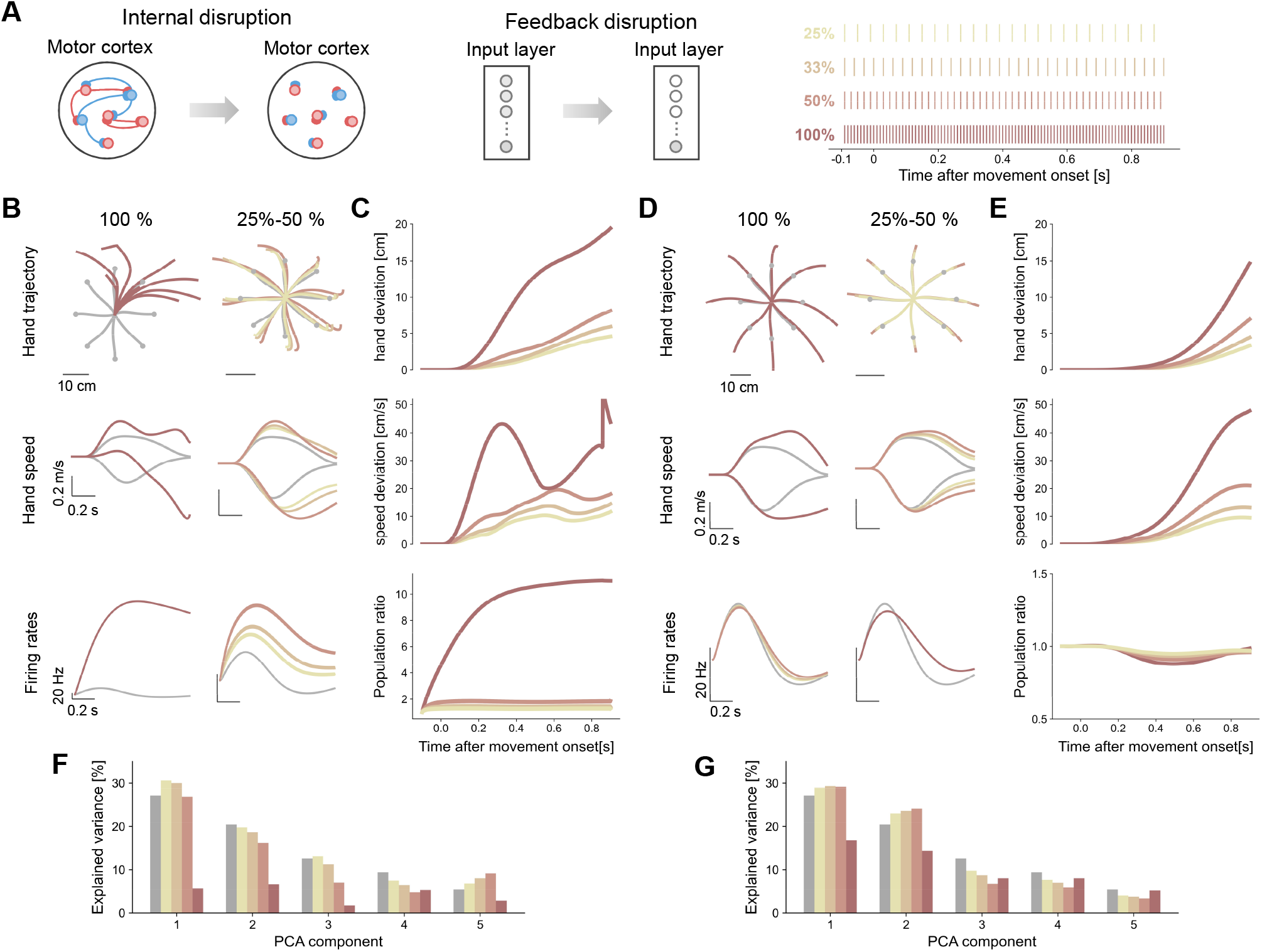
The contribution of internal dynamics and proprioceptive feedback. (A) Illustration of graded disruption. The weights of internal recurrent connections or proprioceptive inputs was set to zero from 25% to 100% timecourse at equal time intervals, respectively. Note, the target input was preserved. (B) The influence of internal dynamics disruption on hand movement and neuronal response. 100% disruption is shown individually, while 25% - 50% are shown together. Top: hand trajectory after disruption compared to the control trial. Middle: x-y hand velocity in condition 1 after disruption compared to the control trial. Bottom: an example neurons’ firing rates in condition 1 after disruption compared to the control trial. (C) The changes of hand movement and neuronal response after internal dynamics disruption. Top: condition-averaged Euclidean distance from the hand position to the control hand position at the same time point for different grades of disruption. Middle: condition-averaged Euclidean distance except for hand velocity. Bottom: population ratio that quantifies the firing rate fluctuation relative to that in control trial (D, E) Same as (B, C) except for proprioceptive feedback disruption. (F) Percentage of variance of the control (grey) and internal-disrupted (yellow to pink) activity after projected onto the space spanned by top ten principal components of control activity. (G) Same as (F) except for feedback-disrupted activity.

To examine the main characteristics of the population response, we identified condition-independent (identical across conditions) and condition-dependent (condition specific) components of the population response during the transition from preparing to executing a reaching (in a 1s window starting before movement onset) via dPCA (Brendel et al., 2011; Kobak et al., 2016). dPCA explains nearly the same amount of variance as PCA on both monkey and model data (top of Figure 3C, E). Condition-independent components capture 54 % variance of monkey C data, 55 % variance of model data (bottom of Figure 3C, E), indicating that the contribution of variance explained by components from real data and model are highly consistent. These results are comparable to the previous finding that the largest component in motor cortical activity is conditionindependent (Kaufman et al., 2016). The two labeled components of model and real data also rank in a similar order. Additionally, the temporal evolution of individual demixed principal components of monkey and model are qualitatively similar (Figure 3D, F). For the conditionindependent components, the first component displays rapid rise before movement onset and slow decay after that, while the second exhibits a bump before movement onset and then decay. The third component of monkey and model are almost identical. For the condition-dependent components, the absolute values of the first two components increase after movement onset and then decrease, while the sign of the third component inverses during movement execution. Additionally, the model’s overall trends of components also closely mirror the trends in the monkey M data. However, the largest component in the monkey M data is conditiondependent (Figure Appendix1B, C), as most of the recorded neurons are heterogeneous.

Moreover, the MC network also display temporal patterns comparable to those observed in previous studies during reaching. First, we find a ring-like structure after applying PCA to firing rates around the go cue (Figure Appendix1B), as reported by Even-Chen et al. (2019). Second, the MC network displays rotational dynamics (Figure (Churchland et al., 2012)A), similar to those in the real data (Figure Appendix2). However, the fitting quality is low in monkey data (Figure Appendix2B). This discrepancy might be due to differences in the analysis window and the identification of movement onset.

To assess the similarity model and real neural response quantitatively., we first applied a modified CCA (Cunningham and Ghahramani, 2015), which finds linear transformations of two datasets to maximize their correlation. The similarity between model and monkey C (M) is 0.88 (0.85); similarity between monkey C and M is 0.87. Moreover, we used Procrustes analysis (PA), which quantifies the degree of alignment between two datasets by *M*^2^ statistic, with a lower *M*^2^ indicating more similar. The *M*^2^ of model and monkey C (M) is 0.23 (0.27); *M*^2^ of monkey C and M is 0.24. These results demonstrate our model is in good agreement with the MC data.

In summary, our MC network model has successfully replicated key features of MC activity in reach task, setting the first stage for further investigations into the cortical pattern generation.

### 3.2. Dissecting the contribution of internal dynamics and proprioceptive feedback

So far we have shown the similarity between model and real data, we ask whether the recurrent architecture resembles the dynamics in MC, or proprioceptive feedback is the key factor of motor cortical dynamics? To investigate the contribution of internal dynamics and proprioceptive feedback when modeling, we built a network without recurrent connectivity (unconnected), and an autonomous network which does not receive proprioceptive feedback. After optimization, the first ten PCs of unconnected network explained too much variance, while the first ten PCs of autonomous network explained too little (Figure Appendix3A). For the quantitative analysis, these networks are less similar to the monkey data compared to the proposed model, where the autonomous network is the least similar to real data (Figure Appendix3B). Moreover, for the dPCA results of unconnected network, there is small amount of CIS and the changes in condition-dependent component lag behind that in the real data (Figure Appendix4). For the dPCA results autonomous network, fluctuations in the condition-dependent component are suppressed. In sum, lack of connectivity and proprioceptive feedback reduces the similarity between the model and the real data. Both intrinsic connection and proprioceptive feedback are necessary to produce brain-like signals.

The structural anatomy of the motor cortex shows that neurons are interconnected, so the recurrent connections cannot be ignored. We then ask does the motor cortical control exhibit a bias towards internal dynamics or proprioceptive feedback? To dissect their contribution, we set internal dynamics (or proprioceptive feedback) to zero while keeping the other component intact. This disruption strategy occurred 25%, 33%, 50%, 100% timecourse at equal time intervals after go cue onset (Figure 4A). Notably, the motor and sensory noise are set to zero as to prevent other interference beyond the disruption of internal dynamics and proprioceptive feedback. We perform the disruption strategy on the baseline model, a simplified version that does not consider feedback delay and noise. Additionally, we find the 100% internal dynamics disruption drives the arm out of control (Figure Appendix5), which cannot be estimated.

100% internal dynamics disruption has a distinct impact on motor output and neuronal activity compared to other grades. It led to significant deviation from the original hand trajectory and irregular changes on hand velocity, suggesting internal dynamics is critical in guiding the goal of the movement (Top and middle of Figure 4B, C). The network activity ‘explodes’ without recurrent connection, reflected in a significant increase in the firing rate of single neuron and ten times activation of population responses (Bottom of Figure 4B, C). In addition, the top ten PCs of the activity in the control explain most of the variance, but little variance of the 100% internal-disrupted activity. These results highlight the dominant role of internal dynamics in movement execution. On the other hand, the influence of other internal dynamics disruption increases with the grade regularly. The greater the disruption, the greater the deviation between the actual and the desire movement direction; there is little effect on the explained variance.

In contrast, there are regularities in how different grades of proprioceptive feedback disruption affect motor output. Proprioceptive feedback disruption resulted in ceaseless hand movement towards the target. The greater the disruption grade, the more difficult it is to terminate the movement (Top and middle of Figure 4D, E). Moreover, proprioceptive feedback disruption had little effect on firing rate (Bottom of Figure 4D, E). The influence of the 100% feedback disruption on the explained variance is less than that of the 100% feedback disruption. These results indicate that proprioceptive feedback may be more involved in fine-tuning motor commands rather than generating them.

To summarize, our results show that internal dynamics play a dominant role in generating cortical patterns, whereas proprioceptive feedback is integral to the fine-tuning of motor output.

### 3.3. Proprioceptive feedback tames sensitivity to initial condition

Initial conditions should tolerate the large neural variability; for example, different initial conditions could result in similar movements (Churchland et al., 2006, 2010). Here, we manipulated the initial conditions into an aberrant state via two strategies: add white Gaussian noise or noise in a specific direction to the initial state (Figure 5A). The strength of the perturbation is quantified by the noise to the initial state with a signal-to-noise ratios (SNRs) of 15, 20, 25 and 30 dB. There are 3200 noisy trials. We define the successful trial as the hand moves within a 4 cm around the target and the norm of velocity at the end of the movement is less than 0.2 within 1 second (Figure 5B, left). We first the compared the movement precision of models with and without proprioceptive feedback.

**Figure 5:**
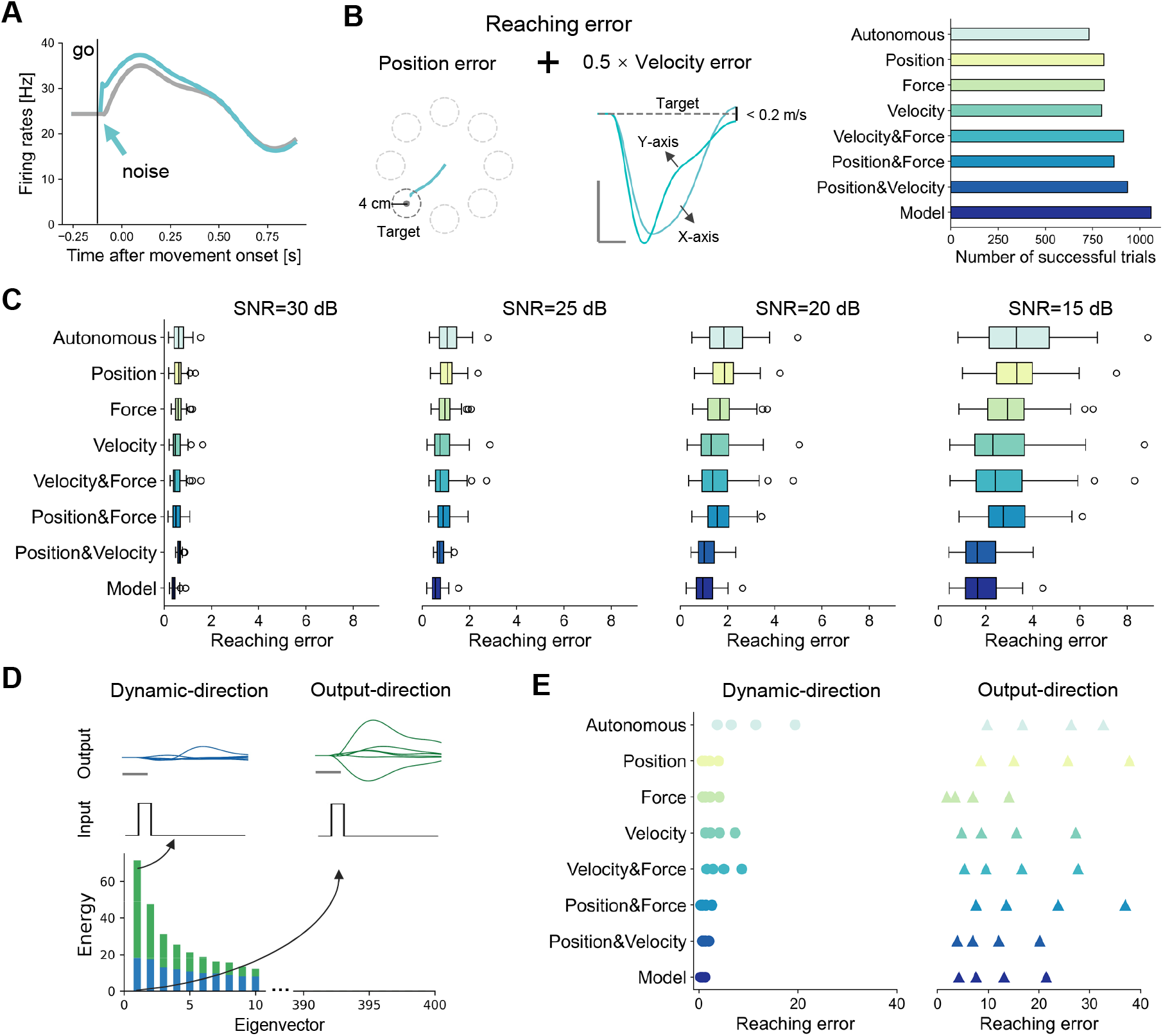
Proprioceptive feedback improves the robustness. (A) Illustration of noise perturbation. Noise is provided to the state of each neuron at go cue onset. Firing rates of an example neuron (cyan) evolve after initial perturbation compared to the control trial (grey). (B) Left: illustration of reaching error which is the sum of position error and 0.5 times velocity error at the end of the movement. Right: number of successful trial of models with different proprioceptive feedback. (C) Box plots of the reaching errors under different SNR conditions of each model. (D) Illustration of the definition of specific noise perturbation, modified from Kao and Hennequin (2019). The eigenvectors of network’s observability Gramians define input sensitivity. Input pulses along the ‘dynamic-direction’ (first blue eigenvector) can trigger large neural responses, while those along the ‘output-direction’ (first green eigenvector) cause large readouts. (E) Reaching errors under different SNR conditions of each model; circles for dynamic-direction perturbation and triangles for output-direction perturbation.

In Gaussian noise perturbation, we find that model completed more successful trial than autonomous network (Figure 5B, right), suggesting that proprioceptive feedback tames the sensitivity to initial condition. The model’s reaching errors is consistently smaller than autonomous model (Figure 5C). Moreover, we perturb the initial conditions in the ‘dynamic-direction’ that evokes the most energy variation of the network dynamics and the ‘output-direction’ that evokes the most energy variation of the readout, respectively. The ‘dynamic-direction’ is defined by the first eigenvector of the observability Gramians whose readout matrix is unit matrix (Figure 5D blue); the ‘output-direction’ is defined by the first eigenvector of the observability Gramians whose readout matrix is the optimized network’s readout matrix (Figure 5D green). The results are consistent with that in Gaussian noise perturbation which further confirm the importance of the proprioceptive feedback (Figure 5E, Appendix6).

In conclusion, model with proprioceptive feedback executed movement with acceptable deviation under different types of perturbed initial conditions. Proprioceptive feedback improves the robustness of the network.

### 3.4. Decoupling of the proprioceptive feedback

The proprioceptive feedback contributes to the robustness of the network. However, it is less clear what the relative importance of components in proprioceptive feedback (hand position and velocity, muscle force). Following the experiment of initial condition perturbation, we built a series of models with different combinations of proprioceptive feedback. We found that decoupling of the proprioceptive feedback from the motor cortex model affects movement precision. The number of successful trial decrease as the number of the proprioceptive feedback types decreases (Figure 5B, right). The reaching error of Gaussian noise perturbation reveals an obvious difference between the model that receives the feedback of hand position and velocity and others (excluding the proposed model) (Figure 5C). Thus, hand position and velocity is relatively important in the proprioceptive feedback. On the other hand, the model receives the feedback of muscle force has smallest error when perturbing in output-direction (Figure 5B, right), suggesting muscle force also plays a role in some specific perturbation conditions.

We further investigated the how different combinations of proprioceptive feedback affect the neural population dynamics. For conciseness, we named the model based on the proprioceptive feedback components, and abbreviated hand position as P, hand velocity as V, muscle force as F. For instance, P&V&F is the model that receives hand position and velocity, muscle force. In the cumulative neural variance, P&V&F and P&V is similar to real data, while other models have smaller amounts of variance; the variance of F is the smallest (Figure 6A). For the quantitative analysis, the order of similarity between simulated and real data is: P&V&F > P&V > V&F > V > P&F > P > F (Figure 6B). There is a clear discrepancy in similarity due to whether models receive hand velocity or not, suggesting hand velocity is relatively important in proprioceptive feedback

**Figure 6:**
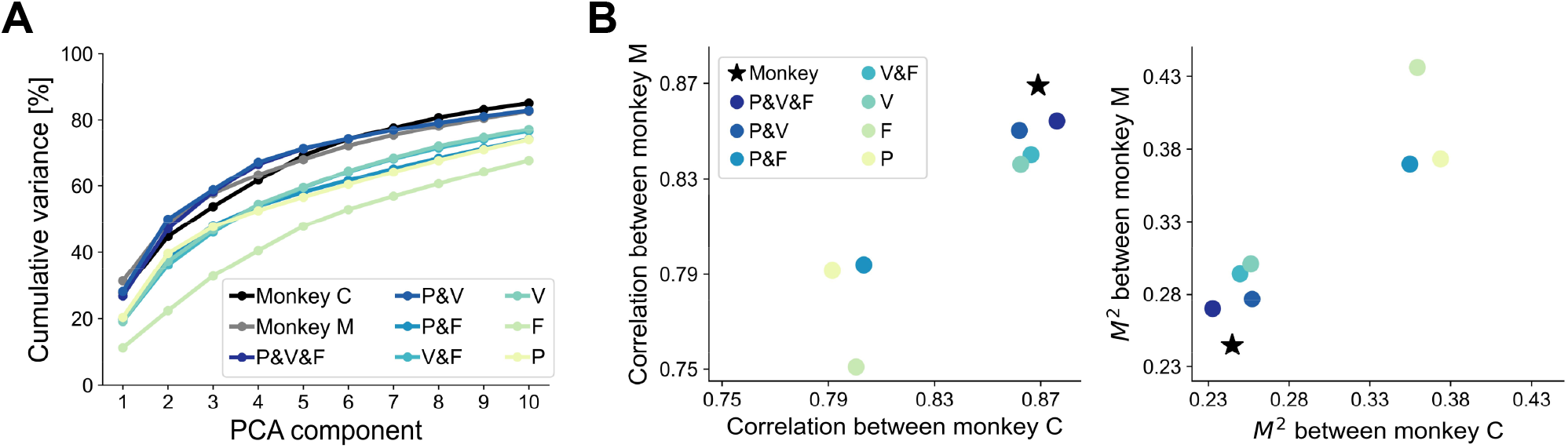
Comparison of simulated and real population responses when decoupling proprioceptive feedback. (A) Cumulative variance explained by first ten PCs of monkey C, monkey M, and models with different combinations of proprioceptive feedback (see text). (B) The similarity between simulated and real data. Left: the canonical correlation between models and monkey M data against the canonical correlation between models and monkey C data. Right: the *M*^2^ between models and monkey M data against the *M*^2^ between models and monkey C data. Note, the similarity between monkey M and monkey C data is shown in black star.

In summary, hand position and velocity is important in improving network’s robustness, and hand velocity had a substantial impact in generating motor cortical dynamics.

### 3.5. Implementation in sensorimotor system

In biological motor system, arm movements are sensed via muscle stretch and stretch velocity rather than the absolute hand position, and possibly muscle force (Vargas et al., 2024). Additionally, proprioceptive feedback is always delayed and noisy Crevecoeur et al. (2016). Arm operates in a relatively unpredictable environment with motor noise. To make our model more biological plausible, we designed a sensorimotor system model, where the motor cortex receive inputs from the proprioceptive system model (Figure 7A). Inspired by Vargas et al. (2024), the proprioceptive system model processes muscle spindle inputs (muscle length and velocity) and efference copy of motor command (muscle activity), predicting hand position, velocity and muscle force. After optimization, the sensorimotor system model also completed the delayed reach task, with similar behavioral performance across trials but trial-to-trial variability within each condition (Figure Appendix1B).

**Figure 7:**
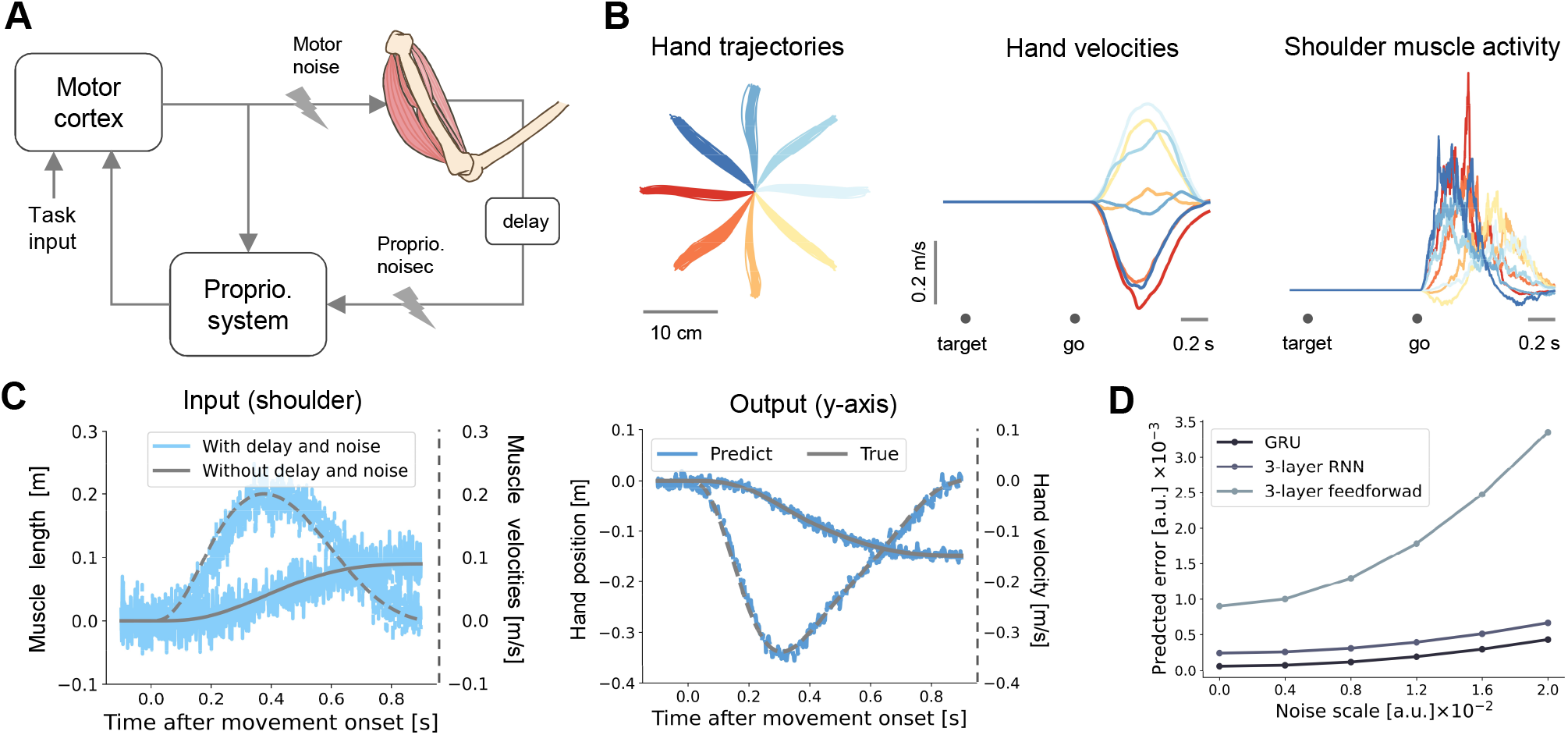
Sensorimotor system model performs delayed-reach task. (A) Illustration of sensorimotor system model. (B) Behaviour performance of the model. Left: Hand trajectories from 40 trials. Middle: X-axis velocities. Right: muscle (shoulder flexor) activity. The squared difference between the real and estimated proprioception under different degree of sensory noise. Black for GRU, dark slate gray for RNN and light slate gray for feedforward network. (C) Inputs and outputs of sensorimotor system model when noise scale is 0.02. Left: The relative length and velocity of shoulder flexor with (sky blue) and without (grey) noise and delay. Right: predicted (blue) and actual (grey) relative hand position and velocity. (D) The squared difference between the real and estimated proprioception under different degree of sensory noise. Black for GRU, dark slate gray for RNN and light slate gray for feedforward network.

Moreover, we found that the sensorimotor system model network achieves high predictive accuracy even under large sensory noise, demonstrating the model’s robustness (Figure 7B). The noise scale *s* represents standard deviation of the Gaussian distribution 𝒩(0, *s*^2^). Moreover, we designed other two models with deeper network layers: a three-layer RNN and a three-layer feedforward network. The results of MSE show that the GRU-based model is the best at estimating the proprioception even under large noise (Figure 7B, D). Therefore, our MC model still works in biologically realistic situations.

## 4. Discussion

It remains an open question how neural activity in MC is generated during movement execution, leading to two two contradictory views that MC is an autonomous (Hennequin et al., 2014; Suresh et al., 2020; Sussillo et al., 2015; Zimnik and Churchland, 2021) or input-driven (Sauerbrei et al., 2020; Kalidindi et al., 2021; Logiaco et al., 2021) dynamical system. Here, we explored the hypothesis that motor cortical activity during movement execution arises from the intersection between internal dynamics within MC and proprioceptive feedback. We built an RNN model of MC that receives proprioceptive feedback (hand position and velocity, muscle force) and controls artificial arm movements. The MC model successfully simulated the neuronal activity in MC during movement execution. The single-neuron activity and population dynamics obtained via various dimensionality reduction techniques are in good qualitative agreement with real MC data. We found that both internal dynamics and proprioceptive feedback contribute to motor cortical dynamics and physical actions.

### 4.1. Dominance of internal dynamics

Internal dynamics predominantly govern cortical pattern generation during movement execution. We observed that disrupting internal dynamics greatly diminished neural activity and made arm movement out of control, compared to the results of disrupting proprioceptive feedback. This suggest that internal dynamics is a key determinant of motor cortical dynamics. Our finding is also consistent with the conclusion in Bachschmid-Romano et al. (2023), who built developed an RNN model of MC under a representational perspective. Furthermore, we can assume that the effect of external inputs is weaker than that of internal dynamics. This can explains why many autonomous dynamical system models generated neural activity similar to MC data, as fast reaching movements are well pre-planned before initiation (Kao et al., 2021; Sussillo et al., 2015).

However, our findings diverge from a previous study (Kalidindi et al., 2021) in which sensory feedback contributed substantially to cortical dynamics. This difference may stem from the different methods of modeling and analysis. First, in the model of Kalidindi et al. (2021), the external input during movement preparation only contains go cue signal and spatial position of the target. This simple combination of feedforward inputs could not capture some remarkable features of preparation, such as various preparatory activity without causing movement and fast preparation(Kaufman et al., 2014; Ames et al., 2014). Theoretical work has provided strong evidence that feedforward input strategy need long time (at least 800 ms) to prepare for a movement (Kao et al., 2021). Thus, the insufficient preparation may led to bias toward the contribution of sensory feedback. Second, we employed a strategy of graded disruption (Sauerbrei et al., 2020) to dissect the contribution of proprioceptive feedback and internal dynamics. It could be rigid to infer from the amplitudes of currents generated from proprioceptive and internal dynamics source, since inputs positioned along the null-dimensions generate no output responses (Kao and Hennequin, 2019). Stavisky et al. (2017) showed that feedback-related motor cortical activity was isolated in null-dimensions that did not affect movement. Third, (Kalidindi et al., 2021) it may be inadequate to prove that network resembles motor cortical dynamics merely by the observation rotational dynamics. Rotational dynamics in the MC is widely observed during various movements, including reaching (Churchland et al., 2012), cycling (Russo et al., 2018), maintaining posture (Kalidindi et al., 2021), and even speaking (Stavisky et al., 2019). Lebedev et al. (2019) argued that rotational structure could be obtained without any realistic properties of motor cortical responses. Therefore, rotational dynamics is probably a fundamental feature of motor cortical activity, but it may not evaluate a model that capture the features of motor cortex sufficiently. Finally, although we also found that the model without recurrent connectivity reflected some motor cortical dynamic characteristics (Kalidindi et al., 2021), we could not claim that a model with these characteristics can simulate MC. There is a need for dependable evaluation index of network simulation.

### 4.2. Proprioceptive feedback in motor control

Proprioceptive feedback could sculpt motor cortical activity. A recent study in the fly shows that motor neurons that control head movement interact with proprioceptive feedback, suggesting that the brain controls movements by a proprioceptive-motor loop (Gorko et al., 2024). We also found proprioceptive feedback plays a modulatory role in motor cortical responses in the disruption experiment.

Proprioceptive feedback is also critical in movement precision. First, MC may control movement velocity through proprioceptive feedback. We observed that disrupting proprioceptive feedback led to continuous, unceasing movement towards the target. This indicates that MC determines when and how to decelerate based on proprioceptive feedback. Also, a study found that the temporal offset in visual feedback extended deceleration duration (McKenna et al., 2017); proprioceptive feedback in our model may have similar effect. Accurate control of movement velocity result in efficient movement execution. The incorporation of sensory feedback in brain-machine interfaces has been shown to enable fast and smooth robotic arm control (Suminski et al., 2010).

Moreover, proprioceptive feedback enhances the network’s robustness. After perturbing the initial conditions with different strategies, the network with proprioceptive feedback still reached the target with much smaller deviation than autonomous model. Decoupling of the proprioceptive feedback from the motor cortex model also led to large deviation from target. Tolerance to the variably of initial conditions is critical in any biologically network, as sensory and motor noise is present in nervous system and different initial conditions can lead to qualitatively similar movements (Churchland et al., 2006). Proprioceptive feedback tames the sensitivity to initial conditions at movement onset.

### 4.3. Bridge between representational and dynamical perspectives

There has been a debate whether MC is better interpreted by representation model and dynamical system model. Our model bridge the gap between representational and dynamical perspectives. We observed an obvious difference in the similarity between simulation and real data when ISN model receive hand velocity feedback or not. This result highlights the contribution of hand velocity feedback to motor cortical activity. First, researches from the view of representation model revealed motor cortical representation of hand position and velocity during reaching movements. Velocity was dominant in motor cortical activity during reaching (Ashe and Georgopoulos, 1994). More than half of neurons in MC are tuned to velocity, and the velocity depth of modulation is significantly greater than the position depth of modulation (Wang et al., 2007). Our results align with the representational perspective that velocity is represented greater than position in MC. On the other hand, our MC model is a dynamical system where neural activity evolves as a result of both the internal dynamics and proprioceptive feedback. Thus, we raise the possibility that MC receives sensory feedback so that its neural activity ‘represents’ motor parameters. The representational and dynamical perspectives do not conflict. Wang et al. (2022) also emphasize that the dynamical systems perspective is not a refutation of the representational perspective but rather a complement or extension.

### 4.4. Proprioception in sensorimotor system

Our MC model captures the essential features of motor cortical dynamics, which could be used to investigate neural basis of motor control. However, the proprioceptive pathway is simplified into a direct feedback of hand motion. To improve the biological plausibility, we built a proprioceptive system model. The key design principles of our model are grounded in the work of Vargas et al. (2024), who found that neurons in proprioceptive system (cuneate nucleus and primary somatosensory cortex) processes muscle spindle inputs, and predicts the position and velocity of the arm, irrespective of the coordinate frameworks. They also suggests that proprioceptive neurons may receive efference copy of motor command might in voluntary movement. In this study, the efference copy of motor command is considered as muscle activity. We considers proprioception is represent in combination of the hand position and velocity, and muscle force in egocentric coordinate from muscle spindle input and muscle activity.

Our sensorimotor system model also reflects the idea of OFC theory, which postulates that MC aims to produce a desired movement and force, taking into account the state of the muscles (Scott, 2004, 2008). According to OFC theory, motor control includes two basic processes. First, an efference copy of motor command is used internally to predict the consequences of movement. These internal predictions are combined with various sensory feedback signals to estimate of the state of the body. Then, the controller (MC) uses this state estimate to adjust motor command. In this work, the proprioceptive system model that receives muscle activity and muscle spindle signals, predicts proprioception. This model play a similar role to a state estimator in OFC theory.

### 4.5. Limitations

There are some limitations in our model. First, to exclude the effect of different starting positions on initial conditions setup, we used the hand position relative to the center point in proprioceptive feedback. However, proprioception represents absolute hand motion (Vargas et al., 2024). Second, the simulation of reaction period is ignored in our model. We assume that internal dynamics and stereotyped condition-independent signal shape the population, while response feedback has a little effect during reaction period. Second, our model need to be extended to more realistic and complex motor tasks to further investigate the contribution of proprioceptive feedback in movement execution. Finally, our model excluded visual feedback, as estimation of hand motion is similar with or without vision (Crevecoeur et al., 2016; Kasuga et al., 2022). In the future, We will investigate the intergration of proprioceptive and visual feedback in motor cortical responses.

In conclusion, our model offers a computational framework to understand the interplay between cortical internal dynamics and proprioceptive feedback during movement execution

## Acknowledgements

This work was funded by the STI 2030—Major Projects (2022ZD0208604), and the National Natural Science Foundation of China (62176151 and 61773259). We are grateful to Guillaume Hennequin and Ta-Chu Kao for modeling instructions; to Juan Gallego and Matthew Perich for sharing the monkey data, and to Jieji Ren and Di Zhu for discussions.

## Data and code availability

The monkey datasets have been deposited on Dryad by Gallego Juan. The code will be available on request.

## A. Appendix

Here we present the computational details of our model and supplementary figures to confirm our results.

### A.1. Thalamocortical circuit model

The dynamics of thalamocortical circuit model can be described as:

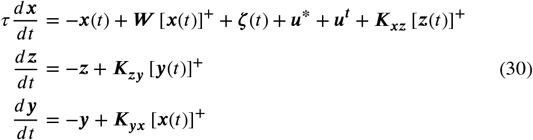

where ***x, y, z*** are the neural activations of MC, MC-layer4 (MC-L4) and thalamus, respectively. []^+^ is the ReLU Activation Function. ***K***_***xz***_ is the MCL4 to MC connectivity matrix, ***K***_***zy***_ is the thalamus to MC-L4 connectivity matrix, ***K***_***yx***_ is the MC to thalamus connectivity matrix. Note, ***K*** = ***K***_***xz***_***K***_***zy***_ ***K***_***yx***_ summarizes these three connectivity matrices around the loop into one feedback gain matrix, which can be calculated from the method described in (Kao et al., 2021). Several constraints were implemented to ensure both biological plausibility and computational feasibility: (1) MC was divided into MC and MC layer 4, and all the connection in the circuit satisfied Dale law. (2) integration dynamics in the thalamus and the MC layer 4 is omitted. (3) muscle activity was set to 0 during movement preparation, bypassing spinal cord modeling for posture control.

#### A.2. Details of arm model

The parameters in force-length-velocity function are: β = 1.55, ω = 0.81, ρ = 2.12, *V*_max_ = −7.39, *c*_*V* 0_ = −3.21, *c*_*V* 1_ = 4.17, *b*_*V*_ = 0.62, *a*_*V* 0_ = −3.12, *a*_*V* 1_ = 4.21, *a*_*V* 2_ = −2.67.

The moment arm matrix *A*, optimal length matrix ***L***^**0**^, and optimal joint angle **θ**^**0**^ are:

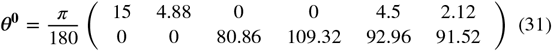

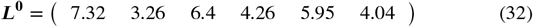

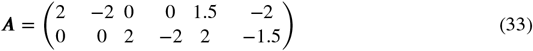

#### A.3. Explanation for the definition of the inputs to the network

We found ***x***(*t*_go_) > 0 generally, thus the dynamics of MC at go cue onset can be linear, [***x***(*t*_go_)]^+^ = ***x***(*t*_go_). Because target is always present after target onset, we assumed that the task input (informing the network where the target is) should contribute to the establishment of initial condition and the following movement execution. The task input in 2.1.2 can be formed as:

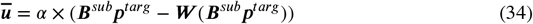

The proprioceptive feedback input in 2.1.2 can be formed as:

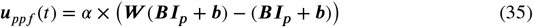

When the hand has reached the target, proprioceptive feedback is:

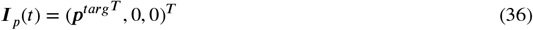

Recall that ***B***^*sub*^ is the first two columns of ***B***. The sum of task input and proprioceptive feedback of hand position is:

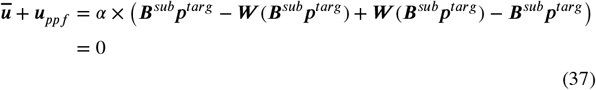

Therefore, the MC network receives the error between current hand position during movement execution.

#### A.4. Ring-like pattern

We first averaged the trial and then averaged firing rate in a 50 ms before go cue, leaving a matrix *X*_*go*_ ∈ *R*^n×*c*^, where n is the number of neurons, c is the number of reach conditions. We preprocessed the data following the procedure in 2.3.1 but we fully normalized each neuron’s firing rate (fr): *f r* = *f r*/*range*(*f r*).We subsequently performed PCA to get *X*_*red*_ ∈ *R*^k×*c*^, where *k* = 2 is the number of principle components and plot the *X*_*red*_ in a plane spanned by top two principle components.

#### A.5. Rotational dynamics

Rotational structure in (Figure Appendix2) is obtained by jPCA, a dynamical variant of principal component analysis (PCA) (Churchland et al., 2012). We first used PCA to reduce the dimensionality of trial-averaged data to *X*_*red*_ ∈ *R*^k×*ct*^, where n is the number of neurons, c is the number of reach conditions, and t is the number of time points, k is the number of principal components. jPCA considers the population responses can be fit into a linear dynamical system 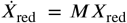, where M is the state-transition matrix. We found the unconstrained *M* that best fits the data *X*, and also identified a constrained skew-symmetric matrix *M*_skew_ that has purely imaginary eigenvalues, in the form of 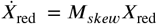. The two leading eigenvectors of the *M*_*skew*_ define the plane where the neural trajectory rotates most strongly.

#### A.6. Models of proprioceptive system

. We modeled proprioceptive system based on GRU, which consists of two gates (Figure Appendix7). The reset gate controls how much of the previous state to remember, while the update gate determines how much of the new state is a copy of the previous state. For the the hidden state of the previous time step, ***h***(*t* − 1), the reset gate ***r***^′^ (*t*) and update gate ***u***(*t*) are computed as follows:

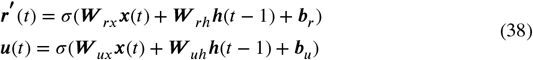

where ***W*** _*rx*_, ***W*** _*ux*_, ***W*** _*rh*_ and ***W*** _*uh*_ are weight parameters.***b***_*r*_ and ***b***_*u*_ are bias. σ is the Sigmoid function.

Then, the candidate hidden state 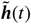 at time t are computed as:

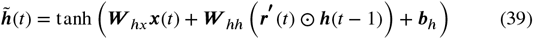

where ***W*** _*hx*_ and ***W*** _*hh*_ are weight parameters. ***b***_*h*_ is bias. ***h*** is bias. ⊙ the Hadamard (elementwise) product operator. The final GRU update equation is defined as:

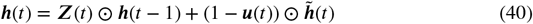

The hidden state is mapped into estimated hand position and velocity, and muscle force:

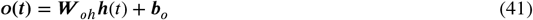

where ***W*** _*oh*_ is weight parameters, and ***b***_*h*_ is bias. There are 200 units in GRU, and all the parameters were sampled from the 𝒩 (0, 0.01)

We also attempted to model proprioceptive system based on other types of artificial neural network. The 3-layer RNN is composed of three RNNs, whose hidden state is referred to as ***h***^**1**^, ***h***^**2**^, ***h***^**3**^. The dynamics of these layers are:

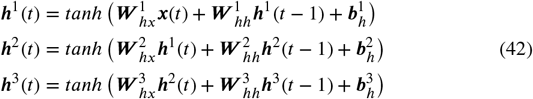

where ***x***(***t***) is the proprioceptive feedback. 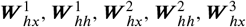, and 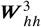 are weight parameters. 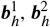 and 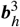 are bias.

On the other hand, the three-layer feedforward network is composed of three linear layers and corrosponding activation function, whose state is referred to as ***z***^1^, ***z***^2^, ***z***^3^. The dynamics of these layers are:

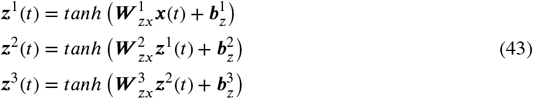

where 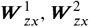 and 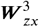 are weight parameters. 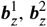 and 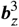 are bias.

**Figure Appendix1:**
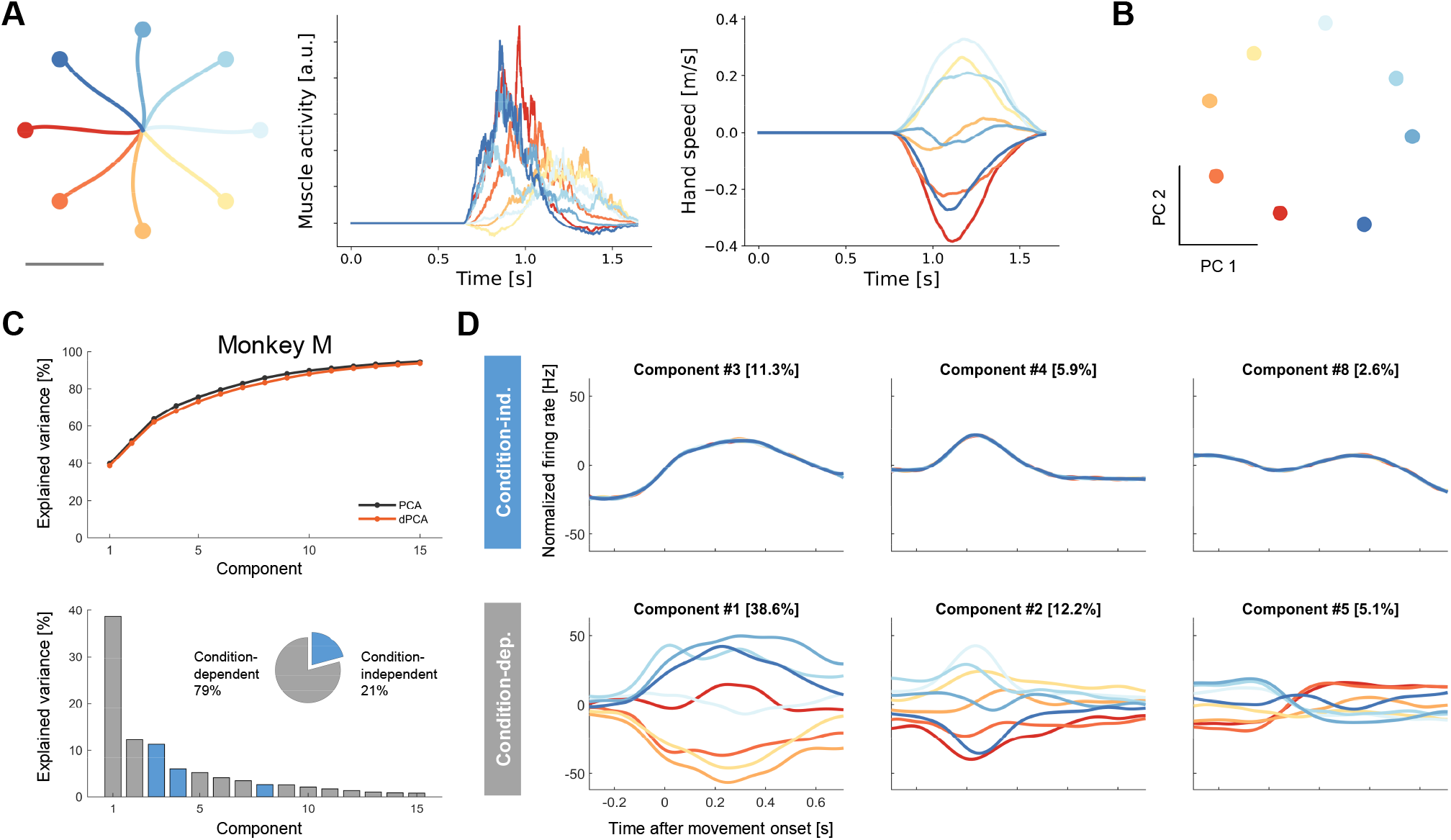
Model and monkey M perform the delayed reach task. (A) Behavioral performance of model within a single trial. From left to right are hand trajectories, shoulder flexor activity, hand velocity (x-axis) in each movmement condition. (B) Ring-like pattern in the plane of top two PCs. (C) Variance captured by dPCA components. Top: cumulative variance explained by dPCA (black) and PCA (orange). Bottom: bars show the variance of the individual demixed principal components. Gray for condition-independent signal, and blue for condition-dependent signal. Pie chart shows the proportions of each signal. (D) Demixed principal components for monkey M data. Top: first three components for condition-independent signal. Bottom: first three components for condition-dependent signal.

**Figure Appendix2:**
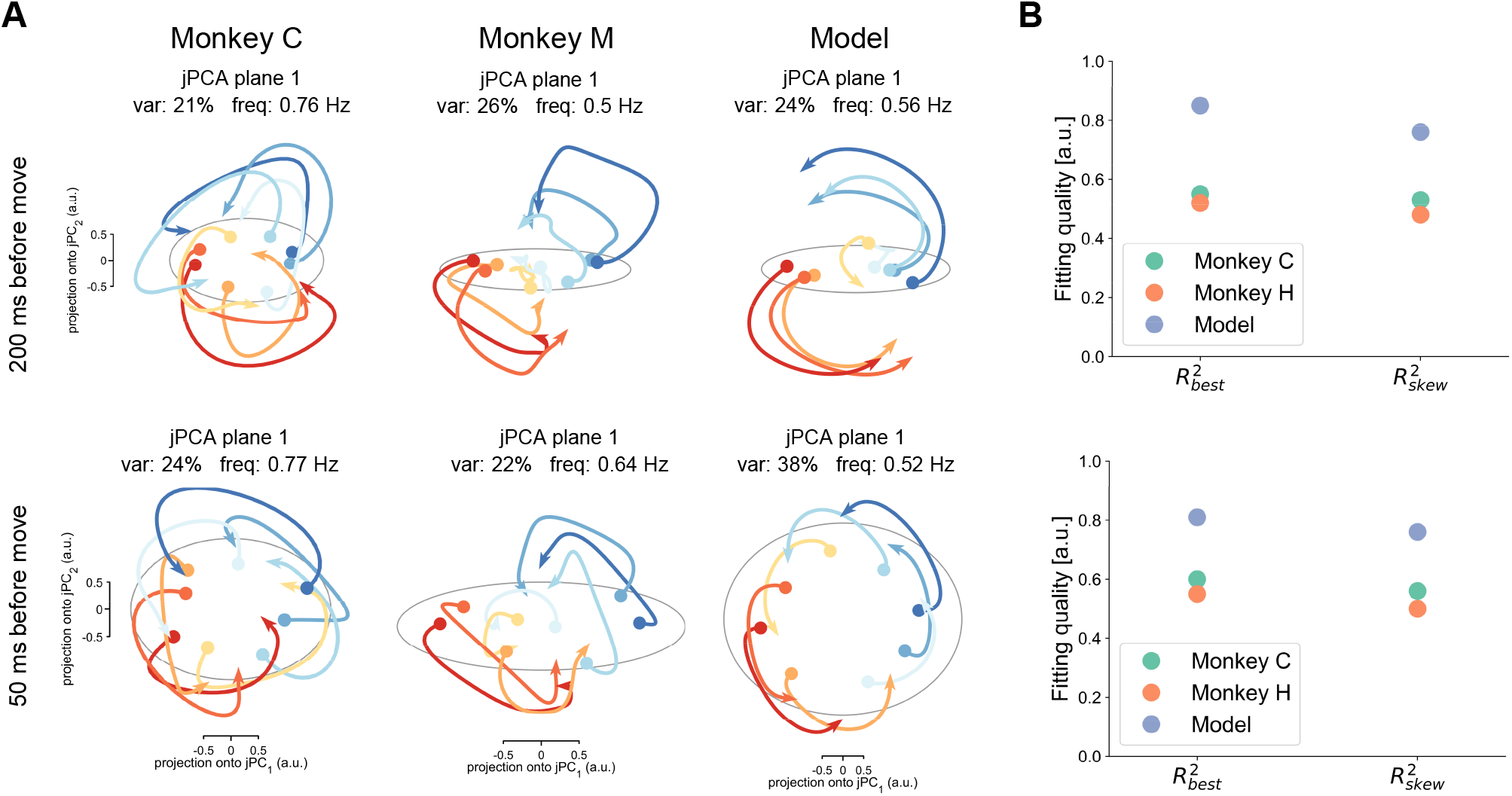
Rotational dynamics of the network and monkeys. (A) Firing rates of monkey C (left), monkey M (middle) and model (right) were projected into top jPC plane. Top, firing rates in a window of 500 ms starting 200 ms before movement onset. Bottom, firing rates in a window of 350 ms starting 50 ms before movement onset. Traces are colored according to the preparatory state projection onto *jP C*_1_. Circle represents initial condition. Grey ellipse shows the range of initial conditions. (B) The fitting qualities for the constrained and unconstrained dynamical systems.

**Figure Appendix3:**
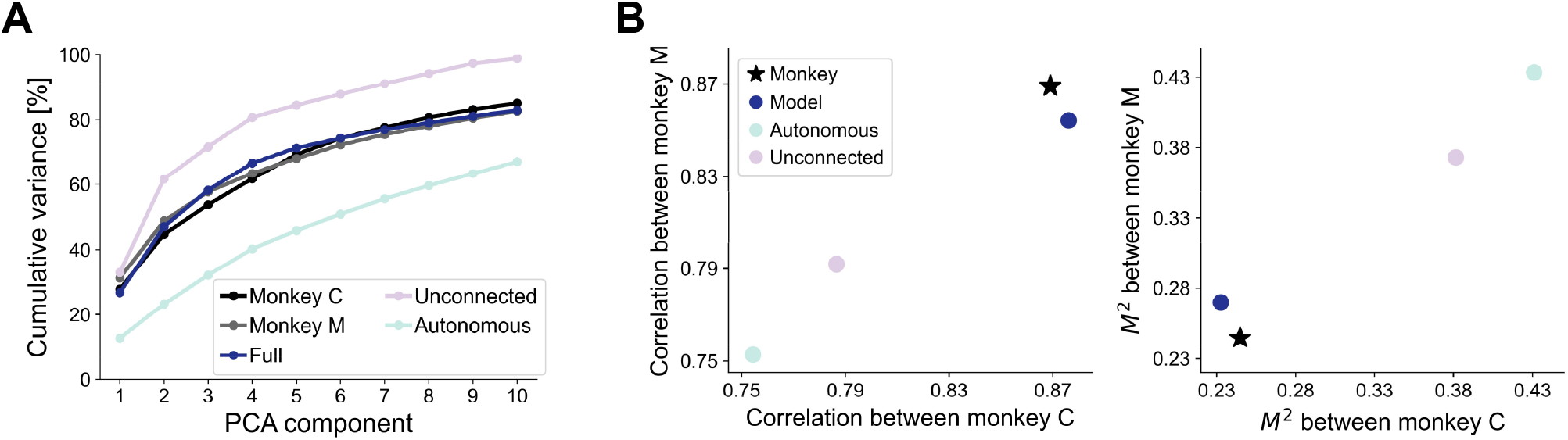
Comparison of simulated and real population responses. (A) Cumulative variance explained by first ten PCs of monkey C, monkey M, the proposed, unconnected and autonomous model. (B) The similarity between simulated and real data. Left: the canonical correlation between models and monkey M data against the canonical correlation between models and monkey C data. Right: the *M*^2^ between models and monkey M data against the *M*^2^ between models and monkey C data. Note, the similarity between monkey M and monkey C data is shown in black star.

**Figure Appendix4:**
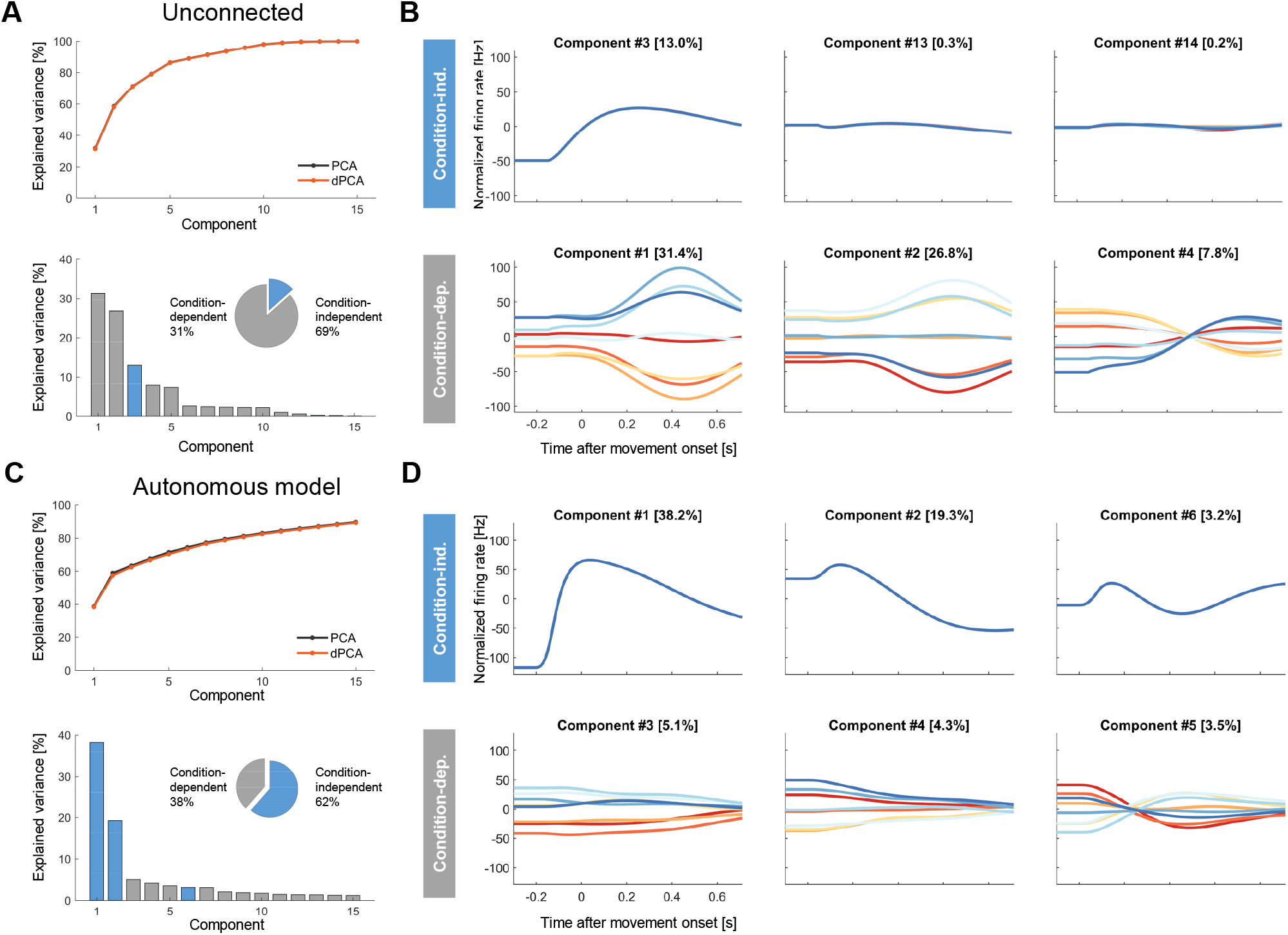
dPCA applied to unconnected and autonomous models. (A) Variance captured by dPCA components for random network. Top: cumulative variance explained by dPCA (black) and PCA (orange). Bottom: bars show the variance of the individual demixed principal components. Gray for condition-independent signal, and blue for condition-dependent signal. Pie chart shows the proportions of each signal. (B) Demixed principal components for random network. Top: first three components for condition-independent signal. Bottom: first three components for condition-dependent signal. (C) Same as (A) except for autonomous network. (D) Same as (B) except for random network.

**Figure Appendix5:**
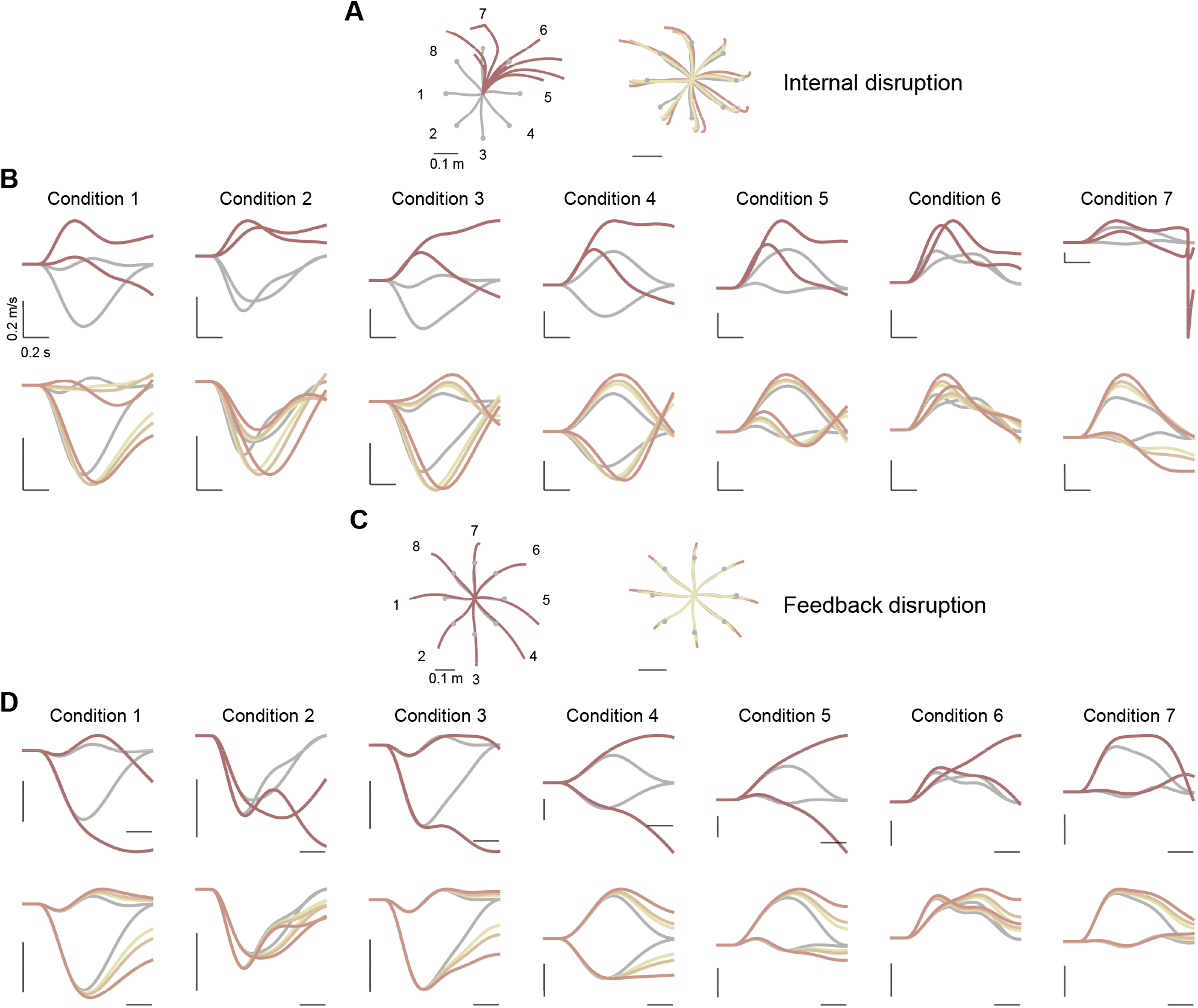
Hand trajectories and velocities in the experiments of internal dynamics and proprioceptive feedback disruption. (A) Hand trajectories for 100% (left) and 25-50%(25, 33 and 50%, right) internal dynamics disruption. Grey bars denote 10 cm. Conditions are numbered counter-clockwise as indicated near the hand trajectories (left). (B) Hand velocities (X and Y axis) for 8 conditions in 100% internal dynamics disruption. (C, D) same as (A, B) except for proprioceptive feedback disruption.

**Figure Appendix6:**
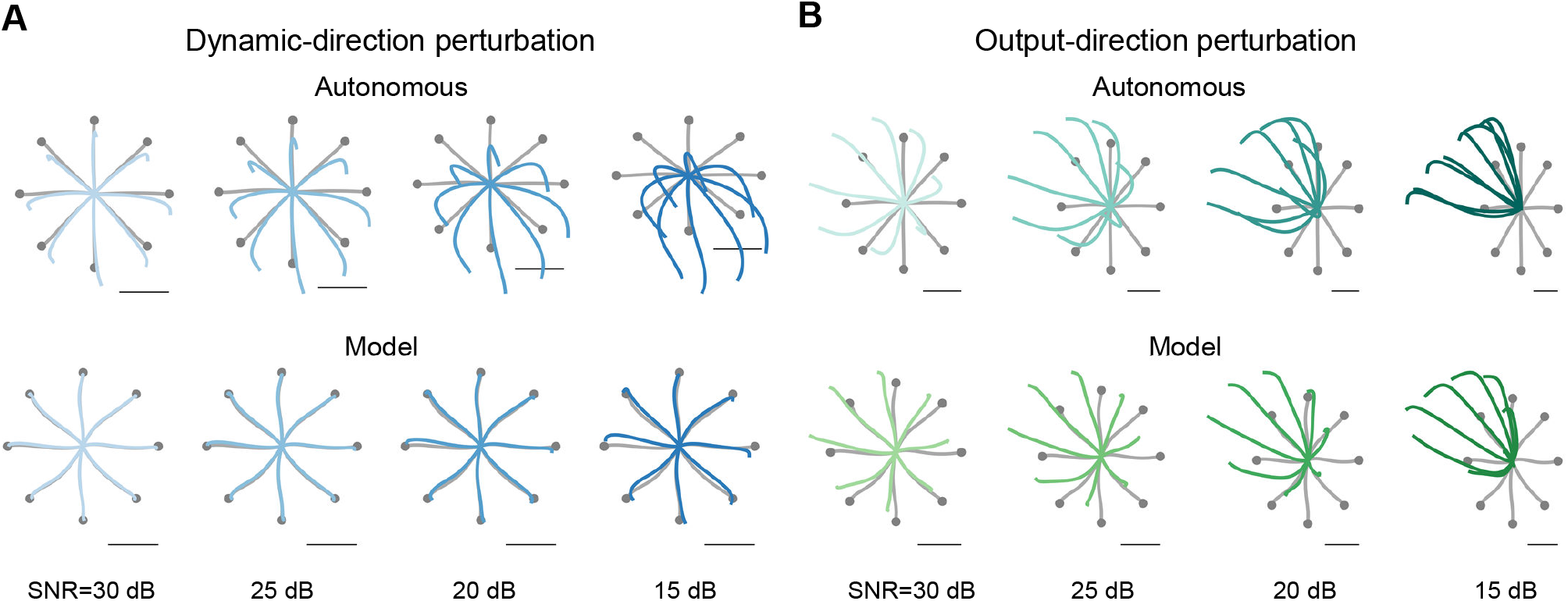
Movement precision when model with and without proprioceptive feedback in initial condition perturbation. (A) Dynamic-direction perturbation. Hand trajectories in the presence (blue) and absence (gray) of noise in the initial condition for one trial. Top: hand trajectories generated by autonomous model. Bottom: hand trajectories generated by proposed model. (B) same as (A) except for output-direction perturbation. As the readout of models are different, hand trajectories in the presence of noise are shown in dark green for autonomous model and green for proposed model.

**Figure Appendix7:**
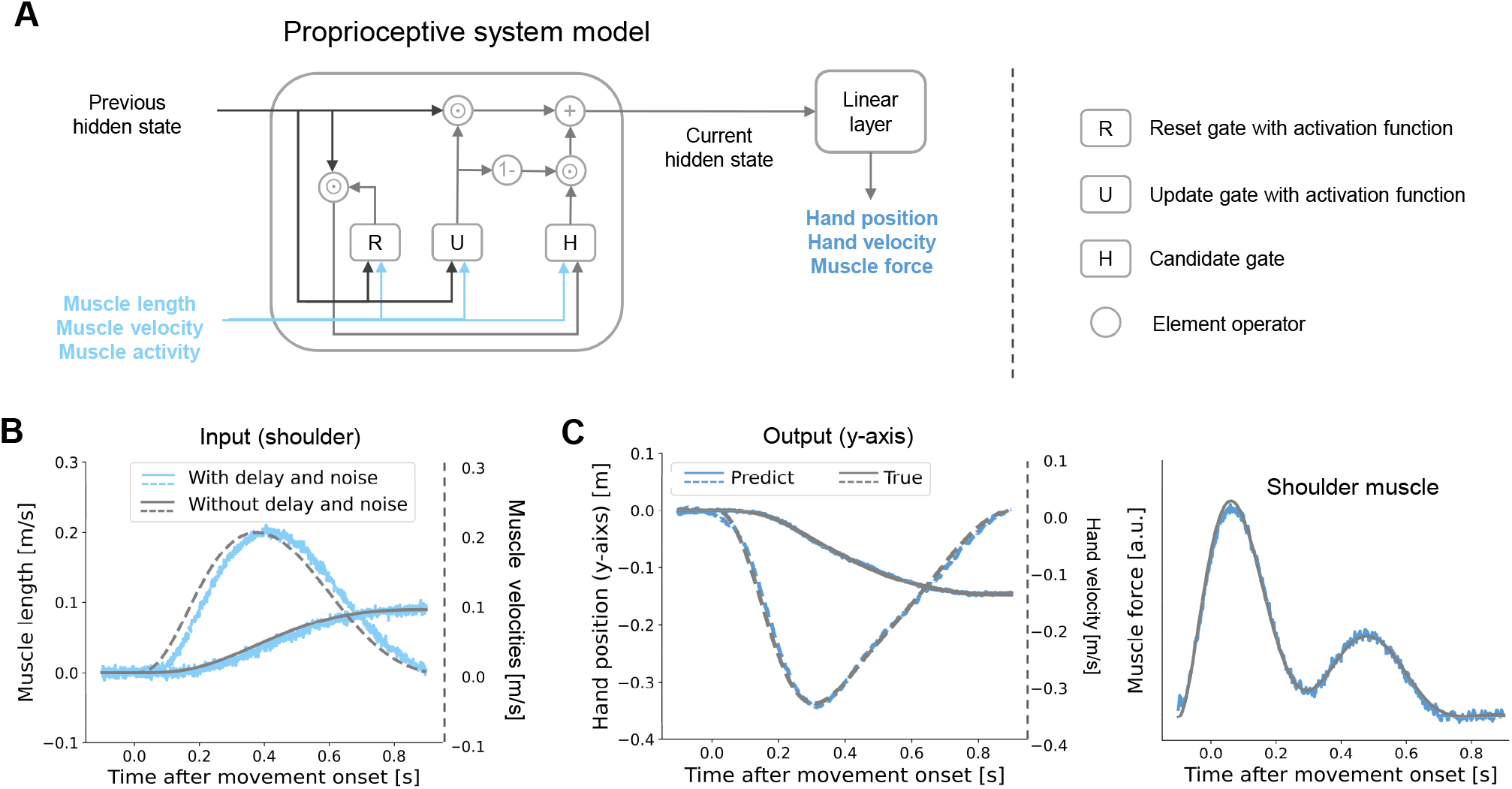
Schematics of a sensory estimation network which receives muscle activity and proprioceptive feedback (muscle length and velocity) with delay and noise, and then predicts hand kinematics and muscle force.

